# T CELLS PROMOTE THE GROWTH OF SMALL-CELL LUNG CARCINOMA VIA AN IL-6/CD74 AXIS

**DOI:** 10.64898/2025.12.19.694894

**Authors:** Maya Baron, Zoé Ginestet, Debadrita Bhattacharya, Myung Chang Lee, Alexandros P. Drainas, Clara L. Poupault, Yoko Nishiga, Alec E. Dallas, Barzin Y. Nabet, Julien Sage

**Author notes:** <u>Authors contributions:</u> M.B. and J.S. designed most of the experiments and interpreted the results; M.B. performed and analyzed most of the experiments with help from Z.G., C.P., and A.E.D, under the supervision of J.S.; M.B., A.P.D., and D.B. performed and analyzed the RNA sequencing data together; Y.N. performed some of the immunostaining analyses; M.C.L. performed data analysis from the IMpower133 trial under the guidance of B.Y.N.; M.B. and J.S. wrote the manuscript and prepared the figures with contributions from all authors.

## Abstract

A central challenge in immuno-oncology is overcoming the limited efficacy and durability of immune checkpoint inhibition (ICI). This is particularly true in small-cell lung cancer (SCLC), where chemotherapy plus ICI is standard of care but rarely curative. Here, we found that T cells are surprisingly both necessary and sufficient for optimal tumor growth in mouse models of SCLC. These pro-tumoral effects are mediated by interleukin-6 (IL-6) production, which induces the pro-survival factor CD74 in SCLC cells. Notably, T cells within human SCLC tumors express IL-6, and low IL-6 signaling correlates with improved survival following chemotherapy plus ICI in patients. Accordingly, IL-6 blockade synergizes with ICI to inhibit SCLC growth *in vivo*. These findings reveal a paradoxical role for T cells in SCLC, uncovering an unexpected T cell–IL-6–CD74 axis that promotes tumor survival, and identify IL-6 as a promising target to help unleash the full potential of immunotherapy in this aggressive cancer.

## Introduction

T cells are a critical component of adaptive immunity and play a central role in orchestrating immune responses against both pathogens and cancer cells. CD8^+^ T cells are central to immune surveillance against cancer, as they possess the ability to directly target and eliminate cancer cells through the recognition of tumor-specific antigens presented by major histocompatibility complex class I (MHC-I) molecules ^1^. CD4^+^ T cells also contribute significantly to responses against cancer cells upon recognition of tumor-associated peptides bound to MHC-II molecules on the surface of antigen-presenting cells (APCs) ^2^. Based on this fundamental knowledge of T cell responses, the development of immunotherapies, such as immune checkpoint inhibition (ICI; e.g., anti-PD-1 or anti-PD-L1), adoptive cellular therapies, and vaccines ^3,4^, has been a paradigm shift for cancer treatment. However, not all cancer patients benefit from these strategies ^5^, and some treated patients suffer from severe side effects, underscoring the need to further investigate the interplay between T cells, cancer cells, and other cells in the tumor microenvironment.

Small-cell lung cancer (SCLC) is one of the most fatal forms of cancer, with a median overall survival of 10-12 months and 5-year survival rates around 10% ^6,7^. SCLC tumors usually occur in heavy smokers, and SCLC cells often have a high tumor mutation burden (TMB) ^8^. While tumor types with a high TMB can respond well to ICI ^9^, only ∼15% of SCLC patients with advanced disease show durable responses in response to ICI ^7,10–14^. One major reason for this relative lack of efficacy is attributed to the low expression of MHC-I molecules on the surface of SCLC cells, which leads to reduced presentation of neoantigens and impedes the recognition and targeting of cancer cells by CD8^+^ T cells ^15–17^. Other possible mechanisms include low infiltration of cytotoxic T cells, a generally immunosuppressive tumor microenvironment, and genetic alterations in SCLC cells that bypass cell death pathways by which T cells ordinarily eliminate cancer cells ^16,18–23^. These mechanisms may be less at play and tumors may be more responsive to ICI in subsets of SCLC patients such as individuals with limited-stage disease ^24^ or whose tumors have a more “inflamed” phenotype ^23,25,26^. SCLC is a neuroendocrine form of lung cancer, and recent evidence suggests that it may also be possible to enhance the anti-tumor effects of ICIs upon DNA damage in neuroendocrine cells or by promoting the differentiation of neuroendocrine SCLC cells towards a less non-neuroendocrine state where cell surface expression of MHC-I molecules is increased ^22,23,27–35^.

Beyond ICI, the development of other types of immunotherapies, including T cells expressing chimeric antigen receptors (CAR T cells) ^36–38^ and bispecific T cell engagers (BiTEs) ^39–41^, further highlights the critical need to investigate the biology of T cells in the SCLC tumor microenvironment. However, the analysis of T cell responses in SCLC is limited by rare surgical resections in this cancer type. It is also challenging to study the interactions between human T cells and SCLC cells in culture or in mouse models. These limitations can be in part circumvented by studying tumors from genetically engineered mouse models of SCLC, including autochthonous tumors that arise in the lungs of mice, as well as allograft models ^22,42–45^.

Using mouse models and data from human tumors, we explored the interaction between T cells and cancer cells in SCLC. In a context where cytotoxic T cells largely fail at killing cancer cells, we uncovered an unexpected pro-cancer effect of T cells on SCLC cells. Specifically, we found that T cells in mouse and human SCLC tumors produce interleukin-6 (IL-6), which promotes the survival of SCLC cells in part by inducing the pro-survival receptor CD74 in the cancer cells. Blockade of IL-6 signaling in SCLC cells in combination with ICI potentiates the effects of these therapies. The identification of pro-cancer T cells in the SCLC microenvironment not only illustrates the complexity of the tumor ecosystem, but also suggests new strategies to enhance the anti-tumor effects of T cell-based immunotherapies.

## RESULTS

### T cells in the SCLC microenvironment promote SCLC growth

We first investigated the role of T cells in a mouse model where SCLC tumors are initiated upon loss of the *Rb1*, *Trp53*, and *Rbl2* tumor suppressors ^46^ (**Supplementary Fig. S1A**). Immunofluorescence analysis of CD3 expression on tumor sections showed a predominant localization for T cells at the tumor borders in this mouse model, with minimal infiltration into the tumor core (**Supplementary Fig. S1B and S1C**), similar to human SCLC tumors ^47–49^. *RPR2* mutant tumors have been shown to have a limited response to ICIs when they grow in the lungs or as subcutaneous allografts ^42,44,50^. Thus, we did not expect T cells to have a major role in tumor growth in the absence of treatments directly or indirectly stimulating their activity in this model, with a possible low basal anti-tumor activity. However, depletion of T cells in *RPR2* mutant mice following injections with anti-CD3 antibody (**Fig. 1A and 1B**, and **Supplementary Fig. S1D**) led to a decrease in lung tumor growth (**Fig. 1C and 1D**). Similarly, when we used the KP1 mouse SCLC cell line (derived from an *RP* mutant tumor ^42,44,50,51^) or the KP11 mouse SCLC cell line (derived from an *RPR2* mutant tumor ^42^), depletion of CD3^+^ T cells led to an inhibition of the growth of subcutaneous allografts in immunocompetent hosts (**Fig. 1E-H** and **Supplementary Fig. S1E-H**). Combined depletion of CD4⁺ and CD8⁺ T cells resulted in similar tumor inhibition phenotypes (**Supplementary Fig. S1I-M**). Collectively, these experiments showed that in a SCLC context where the anti-tumor activity of T cells is limited, these T cells can contribute to tumor progression.

**Figure 1:**
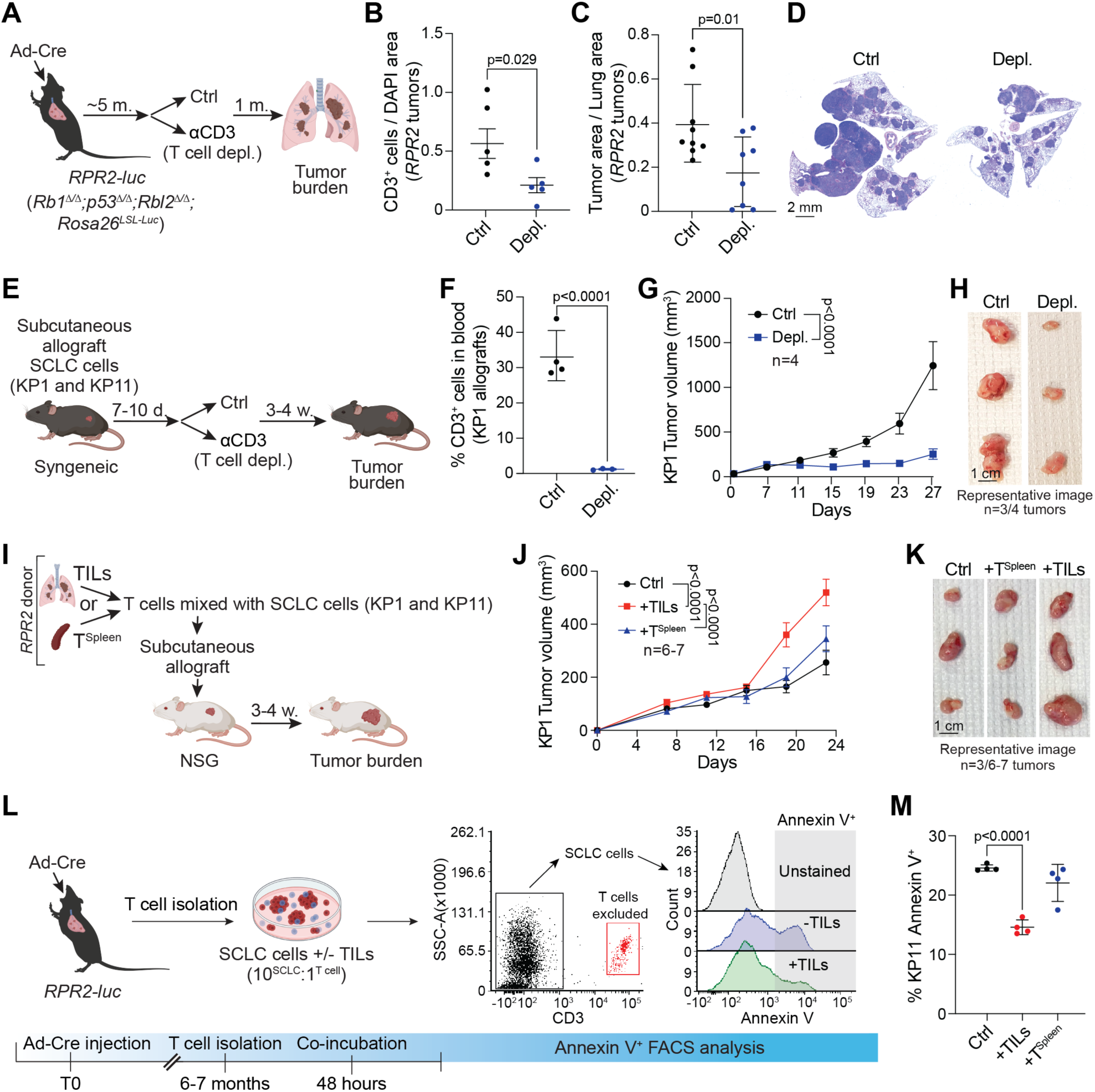
T cells from the SCLC microenvironment promote tumor growth. **A.** Approach used to deplete T cells in the autochthonous *RPR2* model. **B.** Quantification of T cells in the tumor microenvironment of control (Ctrl) and T cell-depleted (Depl.) *RPR2* mutant tumors (n=5), represented as CD3^+^ cells relative to total DAPI area. Statistical significance was assessed using a t-test; error bars represent s.e.m. **C.** Quantification of SCLC tumor area relative to total lung area. Statistical significance was assessed using a t-test; error bars represent s.e.m. **D.** Representative hematoxylin and eosin (H&E) staining of lung sections from control (Ctrl) and T cell-depleted (Depl.) *RPR2* mutant mice (n=8). Scale bar, 2 mm. **E.** Approach used to deplete T cells in mice bearing subcutaneous KP1 SCLC allografts. **F.** Percentage of CD3^+^ T cells in peripheral blood of control (Ctrl) and T cell-depleted (Depl.) mice bearing KP1 SCLC allografts (n=4 mice). Statistical significance was assessed using a t-test; error bars represent s.e.m. **G.** Tumor growth curves of KP1 SCLC allografts (n=4 mice per group, n=1 tumor per mouse, n=1 experiment). Statistical analysis was performed using two-way ANOVA; error bars represent s.e.m. **H.** Representative images of KP1 SCLC allografts at the terminal time point from control (Ctrl) and T cell-depleted (Depl.) mice, as in (G). Scale bar, 1 cm. **I.** Approach used for the T cell transfer experiment. T cells were co-injected with KP1 SCLC cells into NSG mice at a 1:10 ratio (T cells:SCLC cells). **J.** Tumor growth curves of KP1 SCLC allografts in NSG mice injected with SCLC cells alone (Ctrl) or co-injected with splenic T cells (T^Spleen^) or tumor-infiltrating lymphocytes (TILs) (n=6-7 mice per group, n=1 tumor per mouse, n=1 experiment). Statistical analysis was performed using two-way ANOVA; error bars represent s.e.m. **K.** Representative images of KP1 SCLC allografts at the terminal time point as in (J). Scale bar, 1 cm. **L.** Schematic showing the experimental design for co-culture of KP1 SCLC cells with T cells isolated from the spleen (T^Spleen^) or SCLC tumors (TILs). After 48 hours of co-culture, CD3^+^ T cells were excluded from analysis, and SCLC cell apoptosis was assessed using Annexin V staining. **M.** Apoptosis analysis of KP11 SCLC cells cultured alone (Ctrl) or co-cultured with tumor-infiltrating lymphocytes (TILs) or splenic T cells (T^Spleen^) for 48 hours. Apoptosis was analyzed only in CD3-negative cells. Statistical significance was assessed using a t-test; error bars represent standard deviation (n≥3 independent experiments).

To determine whether T cells are not only required but may also be sufficient to promote tumor growth, we injected KP1 or KP11 cells either alone or together with CD3⁺ T cells isolated from the tumor microenvironment (tumor-infiltrating lymphocytes, TILs) or from the spleens of tumor-bearing *RPR2* mutant mice (T^Spleen^ cells) in the flanks of NSG immunodeficient mice, which lack functional T cells. (**Fig. 1I-K** and **Supplementary Fig. S1N-P**). In these experiments, we used an effector-to-target ratio of one T cell to ten SCLC cells, to model the small number of T cells generally observed in or around SCLC tumors. Notably, TILs, but not T^Spleen^ cells, enhanced tumor growth in this assay (**Fig. 1J and 1K** and **Supplementary Fig. S1O and S1P**). These observations supported a specific pro-tumor role for SCLC TILs, even outside of their normal tumor microenvironment in the lungs.

We next examined whether the pro-tumor effect of TILs could be recapitulated *in vitro* by assessing apoptosis and cell-cycle dynamics. TILs and T^Spleen^ cells were freshly isolated from tumor-bearing mice and co-cultured for 48 hours with KP11 or KP1 cells (**Fig. 1L** and **Supplementary Fig. S1Q**). T cells in these assays were used without *ex vivo* activation or stimulation to assess their endogenous activity. Whereas T^Spleen^ cells had no detectable effect, TILs reduced the proportion of apoptotic SCLC cells in these assays (**Fig. 1M** and **Supplementary Fig. S1R**). Cell-cycle analysis revealed no significant differences in SCLC cells co-cultured with either TILs or T^Spleen^ cells (**Supplementary Fig. S1S and S1T**). Altogether, these observations suggested that TILs can promote SCLC growth at least in part by directly enhancing tumor cell survival through the inhibition of apoptosis.

### T cells in the SCLC tumor microenvironment produce IL-6

To begin uncovering the mechanisms underlying the pro-tumor activity of TILs in SCLC, we first sought to identify the T cell population(s) that might drive this effect. We used two complementary approaches: (i) single-cell RNA sequencing (scRNA-seq) to characterize the transcriptional landscape of T cells in SCLC, and (ii) a screen of 48 cytokines to assess potential secreted mediators. In both cases, we compared TILs with T^Spleen^ cells from *RPR2* mutant mice with tumors.

A scRNA-seq analysis of CD3⁺ TILs and T^Spleen^ cells showed overall similar populations, with no subsets uniquely present in TILs (**Fig. 2A** and **Supplementary Fig. S2A and S2B**). TILs were enriched for effector and exhausted CD8⁺ T cells, suggestive of some immune activity against SCLC cells in the lungs of *RPR2* mutant mice with tumors. However, potential pro-tumor subsets, including γδ T cells ^52^, regulatory T cells (Tregs) ^53^, and exhausted T cells ^54^, were detected in both CD3⁺ TILs and T^Spleen^ cells. Thus, this analysis did not provide a direct explanation for the specific pro-tumoral activity of TILs.

**Figure 2:**
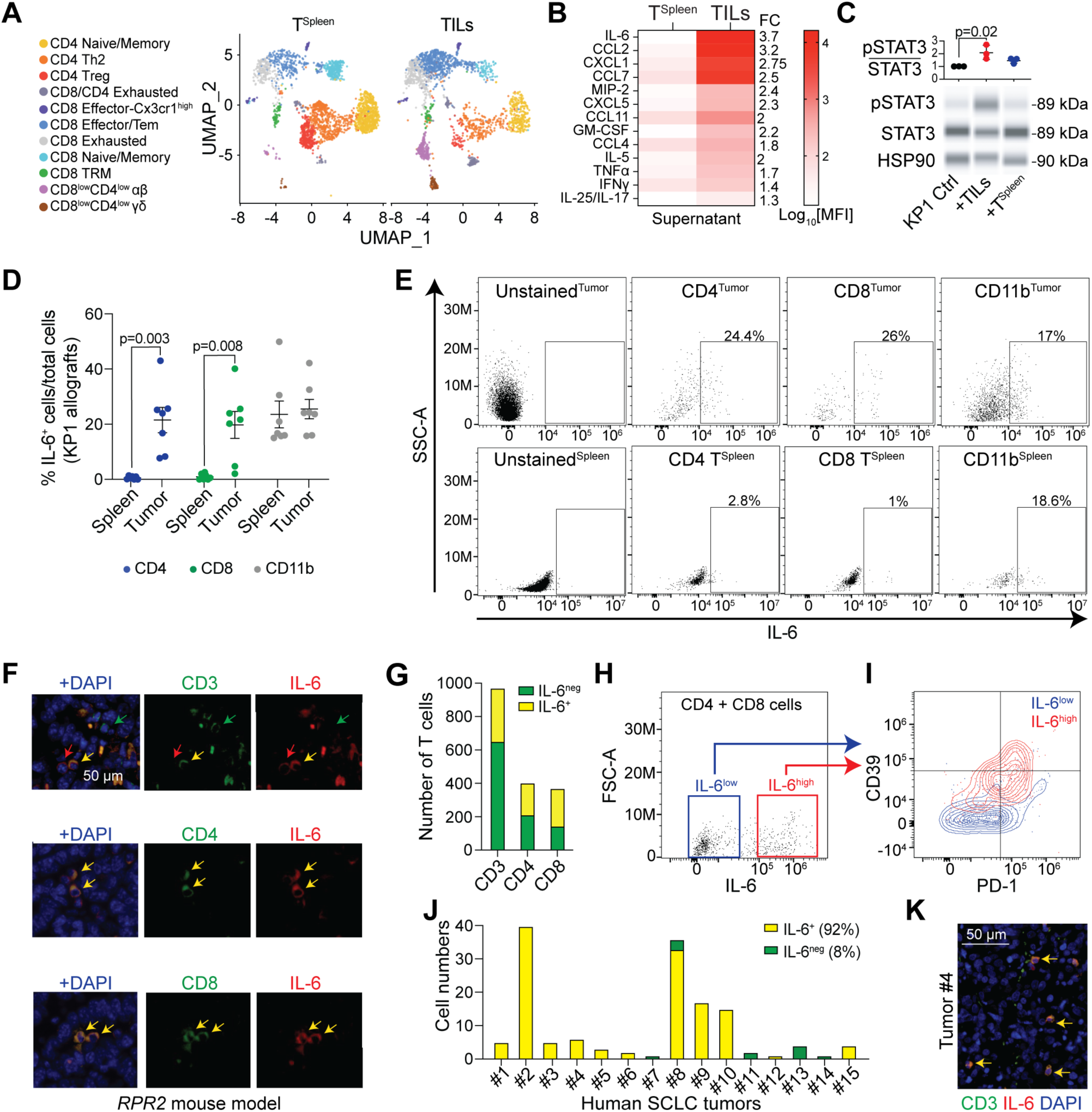
A population of T cells express IL-6 in SCLC tumors. **A.** Uniform Manifold Approximation and Projection (UMAP) plots from single-cell RNA sequencing analysis of tumor-infiltrating lymphocytes (TILs) and splenic T cells (T^Spleen^) isolated from tumor-bearing *RPR2* mice (n=3 mice). Cell clusters are color-coded by cell type. **B.** Heatmap of cytokine levels (measured by Mean Fluorescence Intensity (MFI)) in the supernatant of TILs or T^Spleen^ cells (n=3). **C.** Representative immunoassay and quantification for STAT3 and phosphorylated STAT3 (pSTAT3) levels relative to HSP90 in KP1 mouse SCLC cells grown alone or with TILs or T^Spleen^ cells (n=3). Statistical significance was assessed using a t-test; error bars represent SD. **D.** Percentage of IL-6-expressing cells within total CD4⁺ (blue), CD8⁺ (green), and CD11b⁺ (gray) populations in spleen versus KP1 SCLC allograft tumors (n=7 mice). Statistical significance was determined by a t-test; error bars represent s.e.m. **E.** Representative flow cytometry plots of IL-6 expression in unstained controls and gated CD4⁺, CD8⁺, and CD11b⁺ cells from SCLC tumors (Tumor, top row) and splenic cells (Spleen, bottom row) in KP1 allograft-bearing mice. Each gate indicates the percentage of IL-6⁺ cells. **F.** Representative immunofluorescence images of IL6^+^ TILs from SCLC tumors *RPR2* mutant mice. The IL-6 immunofluorescence signal is shown in red; signal for CD3, CD4, or CD8 is shown in green. DAPI stains the DNA in blue. The red arrow points to an IL-6^+^ CD3^neg^ cell; the yellow arrows point to IL-6^+^ CD3/CD4/CD8^+^ cells; the green arrows point to IL-6^neg^ CD3/CD4/CD8^+^ cells. Scale bar, 50 µm. **G.** Quantification of IL-6^+^ and IL-6^neg^ T cells in the SCLC tumors of the *RPR2* mutant mouse model, as in (F). Data were obtained from n=9 *RPR2* mutant mice with tumors, with at least 15 tumor fields photographed per mouse. **H.** Representative flow cytometry analysis of IL-6 expression in CD4^+^ and CD8^+^ cells isolated from SCLC tumors in *RPR2* mutant mouse. **I.** Representative flow cytometry analysis of PD-1 and CD39 expression in CD4⁺ and CD8⁺ cells, comparing IL-6^high^ (red) versus IL-6^low^ (blue) populations. **J.** Quantification of IL-6^+^ and IL-6^neg^ CD3^+^ T cells in human SCLC tumors by immunofluorescence. CD3^+^ T cells were only detected in 15 samples of a tissue microarray with 80 tumors, each sample representing a distinct patient. **K.** Representative immunofluorescence images of human SCLC sections from tumor #4. The IL-6 signal is shown in red; the signal for CD3 is in green. DAPI stains the DNA in blue. Double-positive cells were detected (yellow arrows). Scale bar, 50 µm.

We next analyzed cytokine secretion from TILs and T^Spleen^ cells. Overall, TILs secreted significantly higher levels of cytokines than T^Spleen^ (**Fig. 2B** and **Supplementary Fig. S2C**), further suggestive of increased activity in tumors. While cytokines characteristic of active T cells, such as TNFα (Tumor Necrosis Factor-alpha) and IFNγ (Interferon-gamma), were elevated in TILs, IL-6 showed the greatest differential expression when compared T^Spleen^ cells (**Fig. 2B**). Consistently, SCLC cells co-cultured with TILs exhibited stronger JAK/STAT (Janus kinase/signal transducer and activator of transcription) pathway activation, evidenced by increased STAT3 phosphorylation, relative to those co-cultured with T^Spleen^ cells (**Fig. 2C** and **Supplementary Fig. S2D**). While high IL-6 levels are often associated with inflammation in the tumor microenvironment, its production is not primarily thought to come from T cells (reviewed in ^55–58^). An analysis of a recently curated scRNA-seq pan-cancer cell atlas ^59^ confirmed these prior observations (**Supplementary Fig. S2E**). Furthermore, little to nothing is known about the effects of IL-6 on SCLC cells.

Since *Il6* mRNA expression was not detectable in the scRNA-seq analysis, we sought to confirm its expression in TILs using flow cytometry. In KP1 and KP11 allograft tumor models, IL-6 was not detected in CD4⁺ or CD8⁺ T^Spleen^ cells, consistent with the cytokine profiling data (**Fig. 2D and 2E** and **Supplementary Fig. S2F-I**). By contrast, analysis of CD4⁺ and CD8⁺ TILs identifies a subset of cells expressing IL-6 in both populations (**Fig. 2D and 2E** and **Supplementary Fig. S2G-I**). The expression of IL-6 in TILs was further validated by immunofluorescence in SCLC primary tumors and liver metastases from *RPR2* mice (**Fig. 2F and 2G** and **Supplementary Fig. S2J**). These results indicate that T cells represent a source of IL-6 in SCLC tumors, alongside other cells in the tumor microenvironment such as macrophages or fibroblasts ^60,61^; in the *RPR2* model, CD11b^+^ myeloid cells also produce IL-6 (**Fig. 2D and 2E** and **Supplementary Fig. S2G-I).**

Further analysis comparing IL-6^high^ and IL-6^low^ expressing CD4⁺ and CD8⁺ cells revealed distinct phenotypic differences (**Fig. 2H and 2I**): IL-6^high^ T cells expressed elevated levels of PD-1 and CD39 (**Fig. 2I**). These markers are commonly associated with T cell inhibition and exhaustion, as well as with dysfunctional and immunosuppressive functions ^62,63^, suggesting that IL-6^high^ T cells represent a tumor-associated subset that potentially contributes to immune suppression within the SCLC tumor microenvironment.

To determine if our findings in mouse tumors extend to human tumors, we performed immunofluorescence on human SCLC tissue sections. T cells were rare in the primary tumor samples analyzed, as expected. However, most of the T cells present expressed detectable levels of IL-6 (**Fig. 2J and 2K** and **Supplementary Fig. S2K**), underscoring the potential pro-tumor role of IL-6-secreting T cells in human SCLC. Notably, we identified IL-6⁺ TILs in colorectal and breast cancer samples (**Supplementary Fig. S2L-O**), indicating that this T cell phenotype extends beyond SCLC (and underscoring that scRNA-seq studies may miss expression of the gene coding for IL-6 ^59^). Thus, IL-6-producing T cells can be a prominent feature of tumors.

### IL-6 signaling blockade inhibits SCLC growth and cooperates with PD-1 blockade

To investigate whether IL-6 derived from TILs can promote SCLC growth, we disrupted IL-6 signaling in SCLC cells. The IL-6 receptor comprises the ligand-binding subunit IL-6Rα and the shared signal-transducing subunit IL-6st (also known as gp130 or CD130). IL-6 signals through two mechanisms: cis-signaling, in which IL-6 binds membrane-bound IL-6Rα and recruits IL6st on the same cell, and trans-signaling, in which IL-6 associates with soluble IL-6Rα or IL-6Rα on neighboring cells to activate IL-6st ^57,58^. Since *IL6ra* deletion would only abolish cis signaling, we instead targeted *Il6st* (**Fig. 3A and 3B** and **Supplementary Fig. S3A and 3B**), which is expected to completely block IL-6-driven signaling in SCLC cells and also prevent compensatory activation by other IL-6 family cytokines, such as leukemia inhibitory factor (LIF), which is also secreted by TILs (**Supplementary Fig. S2C**) and signals via IL-6st. Thus, *Il6st* inactivation provides a more definitive approach to assess tumor cell-intrinsic IL-6 signaling. As expected, *Il6st* knock-out cells were unable to respond to IL-6, as measured by a lack of induction of STAT3 phosphorylation in culture (**Supplementary Fig. S3C and S3D**). Loss of the ability to signal via IL-6st did not affect the growth of mouse SCLC cell populations in culture (**Fig. 3C** and **Supplementary Fig. S3E**), nor did it affect the growth of human SCLC cell lines (DepMap project data, **Supplementary Fig. S3F**).

**Figure 3:**
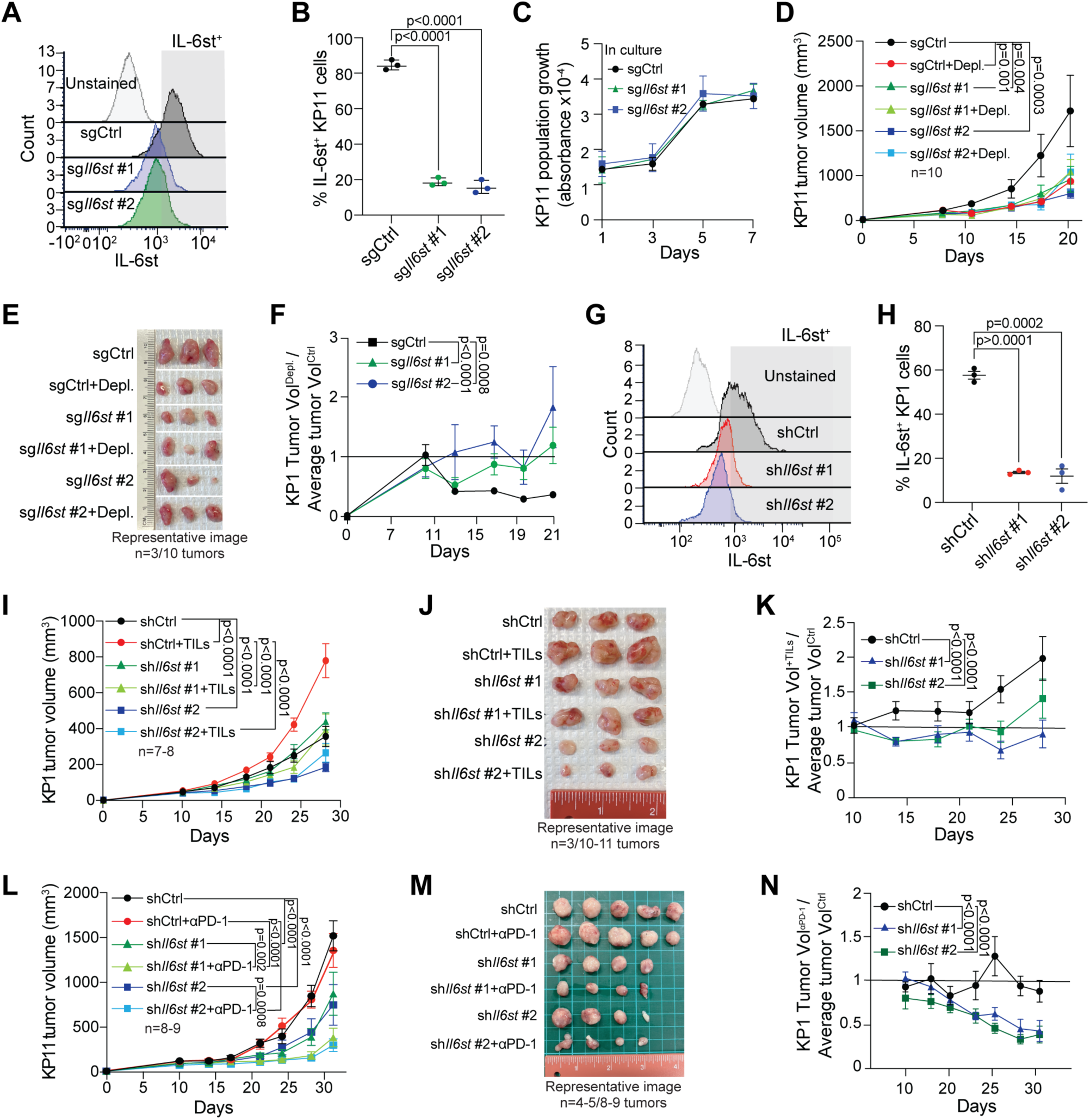
T cells promote tumor growth via the IL-6 receptor in SCLC cells. **A.** Representative flow cytometry analysis for IL-6st levels in control (sgCtrl) or *Il6st* knock-out KP1 cells (sg*Il6st* #1 and #2). Unstained cells provide a negative control for gating. **B.** Quantification of IL-6st expression from (A) (n=3). Statistical significance was determined by a t-test; error bars represent SD. **C.** Cell viability assay (AlamarBlue) of control (sgCtrl) and *Il6st* knock-out (sg*Il6st* #1 and #2) KP1 cells for 7 days (n=3). Statistical analysis was performed using two-way ANOVA; error bars represent SD. **D.** Tumor growth curves of KP1 SCLC tumors, with or without *Il6st* knock-out, in control and T cell-depleted (+Depl.) mice (n=10 mice, n=1 tumors per mouse, n=1 experiment). Statistical analysis was performed using two-way ANOVA; error bars represent s.e.m. **E.** Representative images of KP11 tumors at endpoint, as in (D). **F.** Fold change in tumor size from (D). Tumor volume at each time point was normalized to the mean of control tumors. Ratio >1 indicates an anti-tumor effect of T cells, ratio =1 indicates no effect, and ratio <1 indicates a pro-tumor effect. As expected, ratio <1 for control tumors (T cell depletion slows growth), whereas ratios were closer to or >1 for knock-out tumors. Statistical analysis by two-way ANOVA; error bars represent s.e.m. **G.** Representative flow cytometry analysis for IL-6st levels in control (shCtrl) or IL-6st knock-down KP1 cells. Unstained cells provide a negative control for gating. **H.** Quantification for (G) (n=3). Statistical significance was assessed using a t-test; error bars represent s.e.m. **I.** Tumor growth curves of KP1 allografts injected alone or together with tumor-infiltrating lymphocytes (+TILs), with or without *Il6st* knock-down (n=7-8 mice; 1-2 tumors per mouse; 1 experiment). Statistical analysis by two-way ANOVA; error bars represent s.e.m. **J.** Representative images of KP1 tumors at the terminal time point, as in (I). **K.** Fold change in tumor size from (I). Tumor volume was normalized to the mean of control tumors. Ratio >1 represents a pro-tumor effect of TILs, ratio =1 indicates no effect, and ratio <1 indicates an anti-tumor effect. For control tumors, ratio >1 (TILs promote growth), whereas for knock-down tumors, ratio was closer to 1. Statistical analysis by two-way ANOVA; error bars represent s.e.m. **L.** Tumor growth curves of KP11 allografts left untreated or treated with α-PD-1, with or without IL-6st knock-down (n=8-9 mice; 1-2 tumors per mouse; 1 experiment). Statistical analysis by two-way ANOVA; error bars represent s.e.m. **M.** Representative images of KP11 tumors at the terminal time point, as in (L). **N.** Fold change in tumor size from (L). Tumor volume was normalized to the mean of control tumors. Ratio >1 indicates a pro-tumor effect of T cells, ratio =1 indicates no effect, and ratio <1 indicates an anti-tumor effect. In control tumors, the ratio ∼1 indicates no α-PD-1 effect, whereas in IL-6st knock-down tumors, ratio <1 indicates anti-tumor activity. Statistical analysis by two-way ANOVA; error bars represent s.e.m.

To test whether IL-6 signaling mediates at least in part the pro-tumor functions of T cells *in vivo*, immunocompetent mice were injected with either control or *Il6st* knock-out SCLC cells. The mice were then divided into two groups: one with T cell depletion and one without. To quantify the effect of T cell depletion, we calculated the ratio of tumor size in T cell-depleted versus non-depleted mice. In control tumors, this ratio decreased below 1, indicating that T cell depletion inhibited tumor growth. By contrast, in tumors lacking IL-6 signaling, the depletion ratio remained around or above 1, indicating that T cell depletion had no effect on tumor growth, with only a slight anti-tumor effect observed at the final time point (**Fig. 3D and 3F** and **Supplementary Fig. S3G-K**). This differential response to T cell depletion reveals that IL-6st signaling is required for TILs to exert their pro-tumor functions in the SCLC microenvironment.

As an independent approach to suppress IL-6st signaling in cancer cells and to control for potential immunogenic effects of Cas9 expression, we silenced *Il6st* in SCLC cells using shRNAs (**Fig. 3G and 3H**). We then evaluated whether *Il6st* is required for TIL-mediated tumor promotion by injecting either control shRNA-transduced SCLC cells (shCtrl) or *Il6st* knock-down SCLC cells (*shIl6st* #1 and #2) into the flanks of NSG immunodeficient mice, either alone or together with TILs isolated from tumor-bearing mice. Following a similar methodology, we compared tumor growth with and without TIL addition. The addition of TILs to control SCLC cells accelerated tumor growth, reflected by a ratio >1, as expected. In contrast, no effect was observed when TILs were mixed with *Il6st* knock-down SCLC cells, indicating that IL-6st is required for the pro-tumor effects of TILs in this subcutaneous allograft model (**Fig. 3I-K** and **Supplementary Fig. S3L**)

IL-6 can influence cancer cell sensitivity to PD-1/PD-L1 immune checkpoint blockade ^64,65^. To test whether IL-6st regulates the response of SCLC cells to immunotherapy, we treated mice bearing control or *Il6st* knock-down tumors with anti-PD-1 antibodies. As expected, anti-PD-1 treatment did not alter tumor growth in the control group, as indicated by an anti-PD-1 ratio score of ∼1. By contrast, anti-PD-1 treatment decreased tumor size in *Il6st* knock-down tumors, indicating that blockade of IL-6 signaling sensitizes SCLC cells to PD-1 inhibition (**Fig. 3L-N** and **Supplementary Fig. S3M**). Together, these findings identify IL-6st signaling as a mediator of the pro-tumor effects of TILs in SCLC and a key determinant of tumor sensitivity to anti-PD-1 therapy.

### Tumor-associated T cells promote the survival of SCLC cells via an IL-6/CD74 axis

We next sought to identify the downstream pathways engaged by IL-6 that support SCLC cell survival and tumor growth. To this end, we performed bulk RNA sequencing (RNA-seq) on KP11 SCLC allografts grown in the presence or absence of TILs (**Fig. 4A**). Differential expression analysis revealed 91 genes significantly upregulated and 108 genes downregulated in tumors containing TILs (**Table S1**). To exclude potential contributions of other non-cancer cells in the tumor microenvironment, we performed RNA-seq on KP1 SCLC cells co-cultured with TILs for 48 hours (**Fig. 4A**). Co-culture of SCLC cell with TILs resulted in upregulation of 123 genes and downregulation of 32 genes compared to controls (**Table S2**). Integration of the *in vitro* and *in vivo* datasets revealed only five genes upregulated by TILs in cancer cells in both datasets: *Bhmt*, *Ccdc80*, *Cd74*, *H2-Q7*, and *Pigr* (**Fig. 4B**). To assess the relevance of these candidates in patients, we analyzed bulk RNA-seq data from human SCLC tumors ^66^. Expression of CD3ε (as a proxy for TILs) was correlated only with *CCDC80, CD74*, and *HLA-G* (the human homolog of *H2-Q7*), with the strongest correlation observed for *CD74* (**Fig. 4C** and **Supplementary Fig. S4A-D**). While no well-documented evidence links *Bhmt, Ccdc80, H2-Q7*, or *Pigr* to IL-6 signaling, *CD74* has been reported to be induced by IL-6 in kidney tubular epithelial cells ^67^. To directly assess the role of IL-6 in CD74 regulation in SCLC, we co-cultured TILs with SCLC cells. We observed decreased apoptosis alongside increased CD74 expression in these assays; in addition, blocking IL-6 with a neutralizing antibody not only attenuated the anti-apoptotic effects of TILs but also decreased CD74 induction (**Fig. 4D and 4E** and **Supplementary Fig. S4E and S4F**). To determine whether the presence of TILs may affect CD74 protein expression in SCLC tumors, we analyzed tumor sections by immunofluorescence. We found a positive correlation between TIL density and CD74 expression in both T cell depletion and adoptive transfer experiments (as in **Fig. 1**) (**Fig. 4F and 4G** and **Supplementary Fig. S4G-N**); this association was not due to CD3⁺ cells directly expressing CD74, as the two markers were largely non-overlapping. Consistently, analysis of human SCLC tissue microarrays confirmed a strong correlation between CD74⁺ and CD3⁺ cells (**Fig. 4H and 4I**).

**Figure 4:**
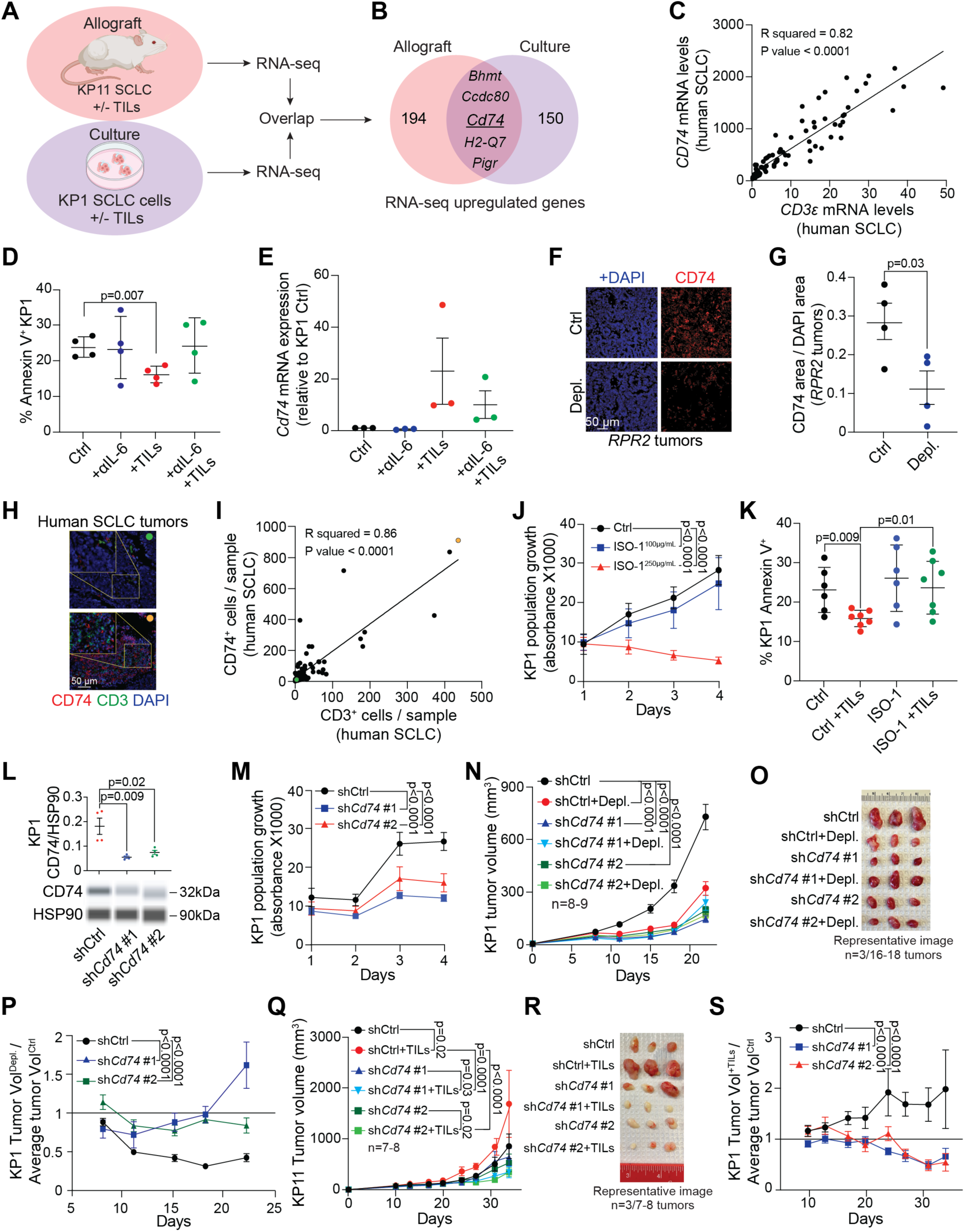
Expression of the pro-survival factor CD74 is induced in SCLC cells by TILs. **A.** Experimental approach. To identify candidate genes mediating the pro-tumor effects of T cells in SCLC, bulk RNA-seq was performed on KP1 cells cultured alone or with TILs (*in vitro*), and on KP11 tumors grown in NSG mice with or without TILs (*in vivo*). **B.** Venn diagram of the number of genes significantly up- and down-regulated in bulk RNA-seq data, as in (A). **C.** Correlation of *CD3ε* (T cells) and *CD74* in RNA-seq data from 81 human SCLC tumors. **D.** Cell death analysis of KP1 SCLC cells treated with IL-6 blocking antibody with or without TILs cells for 48 hours (n=4). Statistical significance was assessed using a t-test; error bars represent SD. **E.** Quantification for *Cd7*4 mRNA levels relative to *Gapdh* in KP1 cells co-culture with TILs, with or without IL-6-blocking antibodies (n=3). Statistical significance was assessed using a t-test; error bars represent s.e.m. **F.** Representative immunofluorescence staining of CD74 (red) in SCLC tumors from control (Ctrl) and T cell-depleted (Depl.) *RPR2* mutant mice. DAPI stains DNA in blue. Scale bar, 50 µm. **G.** Quantification of CD74 staining as in (F), normalized to DAPI staining (n=4 mice). Statistical significance was assessed using a t-test; error bars represent s.e.m. **H.** Correlation of CD3^+^ T cells and CD74^+^ cells from n=110 SCLC samples. Low T cell infiltration (green) and high T cell infiltration (orange) tumors are highlighted. **I.** Immunofluorescence staining of low T cell infiltration (green, right) and high T cell infiltration (orange, left) in human SCLC tumor sections, as in (H). CD3 (green) and CD74 (red). DAPI stains DNA in blue. Scale bar, 50 µm. **J.** Cell viability assay (AlamarBlue) of KP1 SCLC cells treated with ISO-1 (100 µg/mL and 250 µg/mL) or DMSO control (Ctrl) for 4 days (n=3). Statistical analysis was performed using two-way ANOVA; error bars represent s.e.m. **K.** Cell death analysis of KP1 SCLC cells treated with ISO-1 (100 µg/mL) or DMSO control (Ctrl), with or without TILs, for 2 days (n=5-7). Statistical significance was assessed using a t-test; error bars represent SD. **L.** Representatives immunoblot and quantification of CD74 protein levels relative to HSP90 in control (shCtrl) and CD74 knock-down (sh*Cd74* #1 and #2) KP1 cells (n=3). Statistical significance was assessed using a t-test; error bars represent SD. **M.** Cell viability assay (AlamarBlue) of control (shCtrl) and CD74 knock-down KP1 (sh*Cd74* #1 and #2) cells for 4 days (n=3). Statistical analysis was performed using two-way ANOVA; error bars represent s.e.m. **N.** Tumor growth curves of KP1 SCLC allografts in control (shCtrl) and T cell-depleted mice (Depl.), with or without *Cd74* knock-down (n=8-9 mice per group, 2 tumors per mouse, 1 experiment). Statistical analysis was performed using two-way ANOVA; error bars represent s.e.m. **O.** Representative images of KP1 SCLC allografts (terminal time point) as in (N). **P.** Fold change in tumor size as in (N). A depletion ratio score was calculated for each time point by dividing tumor volume by the mean of control tumors. Ratio >1 indicates an anti-tumor effect of T cells, ratio = 1 indicates no effect, and ratio <1 indicates a pro-tumor effect of T cells. In controls the ratio decreased (<1), whereas in CD74 knock-down tumors it remained ∼1. Statistical analysis was performed using two-way ANOVA; error bars represent s.e.m. **Q.** Tumor growth curves of control KP11 SCLC allografts and allografts co-injected with TILs, with or without *Cd74* knock-down (n=7-8 mice per condition, 1-2 tumors per mouse, 1 experiment). Statistical analysis was performed using two-way ANOVA; error bars represent s.e.m. **R.** Representative images of KP11 SCLC allografts (terminal time point), as in (Q). **S.** Fold change in tumor size as in (Q) calculated at each time point by dividing tumor volume by the mean of control tumors. Ratio >1 indicates a pro-tumor effect of T cells, ratio = 1 indicates no effect, and ratio <1 indicates an anti-tumor effect of T cells. A pro-tumor effect was observed in control tumors (ratio >1), whereas no effect was observed in CD74 knock-down tumors (ratio ∼1). Statistical analysis was performed using two-way ANOVA; error bars represent s.e.m.

CD74 is primarily known for its role as an MHC class II chaperone ^68^, but evidence from several studies indicates that CD74 expressed by cancer cells also functions as a pro-survival receptor independent of its MHC class II role ^69–71^. CD74 can promote cancer cell survival via several pathways, including AKT signaling ^72^, a pathway known to be critical in promoting SCLC cell survival ^73^. In co-culture assays with SCLC cells and either TILs or T^Spleen^ cells, we observed activation of AKT, as indicated by an increased ratio of phosphorylated AKT to total AKT, only in SCLC cells co-cultured with TILs (**Supplementary Fig. S4O**).

Based on this evidence, we investigated whether CD74 mediates the pro-tumor effects of TILs in SCLC. Signaling downstream of CD74 is induced by binding to its ligand, macrophage migration inhibitory factor (MIF) ^74^. SCLC cells, both *in vitro* and in allograft tumors, express MIF, providing a source of ligand for CD74 signaling (**Supplementary Fig. S4P and 4Q**). Treatment of SCLC cells in culture with ISO-1, a MIF inhibitor ^75^, led to decreased population growth in culture (**Fig. 4J** and **Supplementary Fig. S4R**), suggesting a basal pro-survival role for the MIF-CD74 axis in SCLC cells. Importantly, short-term treatment with ISO-1 blocked the pro-survival effects of TILs on KP1 SCLC cells in the co-culture assay (**Fig. 4K** and **Supplementary Fig. S4S**). Similar to ISO-1 treatment, knock-down of *Cd74* in SCLC cells led to decreased survival and an inhibition of population growth in culture (**Fig. 4L and 4M** and **Supplementary Fig. S5A and S5B**). In SCLC allografts, *Cd74* knock-down and T cell depletion each similarly inhibited tumor growth, and their combination resulted in no additional tumor inhibition (**Fig. 4N and 4O** and **Supplementary Fig. S5C-G**), further supporting a pro-survival role for CD74 in SCLC cells downstream of interactions with T cells. Consistently, the tumor-size ratio comparing control tumors with T cell-depleted, *Cd74* knock-down tumors remained around or above 1, indicating that T cells had little to no effect (**Fig. 4P** and **Supplementary Fig. S5F**). In reciprocal experiments, co-injection of TILs with control SCLC cells accelerated tumor growth (+TILs ratio > 1), but addition of TILs to CD74 knock-down tumors failed to promote growth (+TILs ratio < 1), indicating that CD74 is essential for T cell-mediated tumor promotion (**Fig. 4Q-S** and **Supplementary Fig. S5H-L**). Collectively, these findings establish a mechanistic axis in which TIL-derived IL-6 induces CD74 in SCLC cells, driving pro-survival signaling and tumor promotion.

### IL-6 blockade cooperates with PD-1 blockade to inhibit SCLC growth

Targeting IL-6st or CD74 in SCLC may be possible in the future but these molecules are not currently targetable in patients. Therefore, we tested whether selective IL-6 blockade, in combination with PD-1 checkpoint inhibition, could provide a clinically relevant strategy. In the KP1 and KP11 allograft models, IL-6 blockade alone or PD-1 blockade alone had minimal to no effect on tumor growth; in contrast, the combination of IL-6 and PD-1 blockade significantly suppressed tumor progression (**Fig. 5A-D** and **Supplementary Fig. S6A-C**), similar to the *Il6st* knock-down (**Fig. 3L-N** and **Supplementary Fig. 3M**).

**Figure 5:**
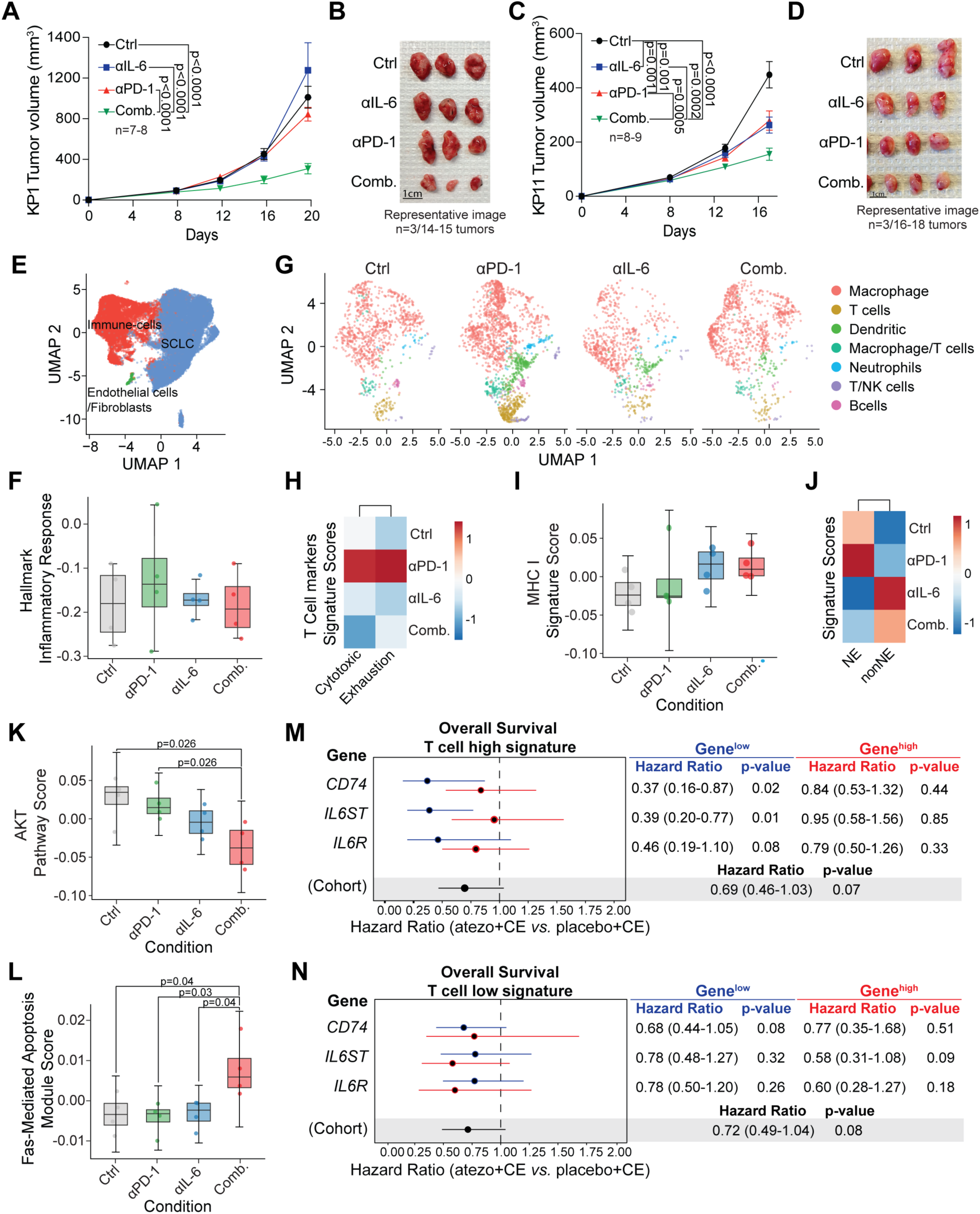
Combining IL-6 and PD-1 blockade inhibits SCLC growth. **A.** Tumor growth curves of KP1 SCLC tumors treated with IL-6 blockade, PD-1 blockade, or the combination of IL-6 and PD-1 blockade (n=7-8, 1-2 tumors injected per mouse, 1 experiment). Statistical analysis was performed using two-way ANOVA; error bars represent s.e.m. **B.** Representative images of KP1 tumors from (A). **C.** Tumor growth curves of KP11 SCLC tumors treated with IL-6 blockade, PD-1 blockade, or the combination of IL-6 and PD-1 blockade (n=8-9, 1-2 tumors injected per mouse, 1 experiment). **D.** Representative images of KP11 tumors from (C). **E.** Uniform Manifold Approximation and Projection (UMAP) plots for the scRNA-seq analysis of KP11 allografts (n=4 per treatment condition). Cell clusters are color-coded by cell populations. **F.** UMAP plots of scRNA-seq data for immune cells (CD45^+^) from KP11 allografts (n=4), untreated or treated with PD-1 blockade, IL-6 blockade, or their combination. Clusters are color-coded by cell population. **G.** Box plot of Hallmark inflammatory response gene signature scores across four experimental conditions: untreated, PD-1 blockade, IL-6 blockade, and IL-the combination (n=4). Statistical analysis by t-test; error bars represent SD. **H.** Heatmap showing cytotoxic and exhausted gene signatures in T cells in the four experimental conditions as in (G). **I.** Box plot for an MHC I signature score in SCLC cells as in (G). Statistical analysis by t-test; error bars represent SD. **J.** Heatmap of neuroendocrine (NE) and non-neuroendocrine (nonNE) gene signatures in SCLC cells as in (G). **K.** Box plot for AKT pathway scores in SCLC cells as in (G). Statistical analysis by t-test; error bars represent SD. **L.** Box plot for FAS-mediated apoptosis gene signature scores in SCLC cells across as in (G). Statistical analysis was performed using a t-test; error bars represent SD. **M.** Forest plots showing overall survival (OS) hazard ratios with 95% confidence intervals (CI) for atezolizumab plus carboplatin/etoposide (atezo+CE) vs placebo plus CE (placebo+CE) in the T signature > median cohort. Low gene expression (≤ median) shown in blue (Gene^low^); high gene expression (> median) shown in red (Gene^high^). “Cohort” refers to T signature > median without further subsetting. **N.** As in (M) for T signature ≤ median and gene expression.

To investigate possible mechanisms underlying the therapeutic effect of the combination treatment, we performed scRNA-seq on untreated tumors, and tumors treated with anti-PD-1, anti-IL-6, or the combination, analyzing both immune cells and SCLC cells (**Fig. 5E**). IL-6 signaling can suppress T cell function and promote tumor immune evasion ^76^. Moreover, IL-6 blockade has been reported to enhance anti-tumor immune responses ^77–79^. Based on these observations, we first checked if the anti-IL-6 treatment affected immune infiltration and activity, thereby augmenting anti-PD-1 therapy. Overall, however, we did not observe significant changes in hallmark inflammatory response in tumors treated with anti-IL-6 (**Fig. 5F**). Quantification of TILs showed no increase in TILs frequency following anti-IL-6 treatment alone (6%) or in combination with anti-PD-1 (5%), compared with control (6%) or anti-PD-1 (15%) (**Fig. 5G** and **Supplementary Fig. S6D**). Moreover, TILs in the anti-IL-6 group of tumors exhibited reduced cytotoxic and exhaustion signatures (**Fig. 5H**), suggesting limited functional activation. Finally, no significant upregulation of MHC I was detected in SCLC cells from treated tumors (**Fig. 5I**), indicating that increased susceptibility to T cell cytotoxicity is unlikely to explain the observed therapeutic benefit. Together, these findings suggest that enhanced T cell activity is not the primary driver of tumor inhibition by anti-IL-6 in SCLC.

Given prior evidence that SCLC tumors with non-neuroendocrine (non-NE) features are more sensitive to immunotherapy ^26,80^, we next evaluated transcriptional programs in SCLC cells, using the KP11 model (**Supplementary Fig. S6E**). Anti-IL-6 treatment was associated with some reduced ASCL1 target gene expression (NE phenotype) and some increased HES1 target gene expression (non-NE phenotype), but these changes did not reach statistical significance (**Fig. 5J**). This reprogramming was similar in the combination therapy, in which tumor inhibition is more pronounced, suggesting that the slight shift towards a less NE state is insufficient to suppress SCLC progression.

We next examined whether CD74 pro-survival signaling may contribute to the therapeutic response. Whereas NF-κB signaling remained unchanged across conditions (**Fig. S6F**), AKT pathway activation was significantly reduced only in the combination treatment compared with control or anti-PD-1 alone (**Fig. 5K**), consistent with our prior observation in SCLC cells co-cultured with T cells (**Supplementary Fig. S4O**). Finally, Fas signaling, which is negatively regulated by CD74 pro-survival pathways ^72^, was upregulated exclusively in the combination group (**Fig. 5L**), suggesting that effective tumor regression requires both inhibition of cancer cell-intrinsic survival signaling (via IL-6 blockade) and enhanced T cell activity (via PD-1 blockade).

We next evaluated the clinical relevance of the IL-6/CD74 axis in SCLC patients. The IMpower133 clinical trial investigated the benefits of the PD-L1-blocking antibody atezolizumab plus carboplatin and etoposide (atezolizumab+CE) in SCLC, compared to placebo plus carboplatin and etoposide (placebo+CE) ^7,81^; RNA-seq data are available from this study and can be used to investigate the survival of subgroups of SCLC patients ^23,26^. In tumors with higher T cell infiltration (estimated using a T cell gene expression signature), *CD74*-low or *IL6ST*-low status (in blue), rather than *CD74*-high or *IL6ST*-high status (in red), was significantly associated with improved progression-free survival (PFS) and overall survival (OS) in the atezolizumab+CE group compared to the placebo+CE group (**Fig. 5M** and **Supplementary Fig. S6G**). This effect was dampened in tumors with low T cell infiltration, where atezolizumab+CE was associated with benefit compared to placebo+CE, irrespective of *CD74, IL6ST*, or *IL6R* mRNA expression levels (**Fig. 5N** and **Supplementary Fig. S6H**). These observations further support a model in which reducing IL-6/CD74 signaling may enhance the benefit of immunotherapies in SCLC patients with baseline lymphocyte infiltration.

## Discussion

In this study, we investigated the functional interactions between cancer cells and T cells in a tumor context where the cytotoxic effects of T cells on cancer cells are minimal. Under these conditions, we found that T cells can be co-opted by cancer cells to function as a supportive stroma that promotes tumor growth. These findings support a model in which T cells exhibit concurrent pro- and anti-tumor activities. In cancers like SCLC, where T cell-mediated tumor cytotoxicity is limited, targeting the pro-tumor functions of T cells may help unmask their anti-tumor potential and aid in controlling tumor growth.

SCLC is typically considered an “immune-cold” cancer type, with most tumors showing low lymphocyte infiltration, poor antigen presentation, and robust immune evasion mechanisms. Still, some tumors have more inflamed phenotypes and respond more effectively to ICI, likely via the activation of cytotoxic T cells ^23,26,81^. A major focus in the field has thus been to find ways to further enhance the anti-tumor effects of T cells. In contrast, our work has focuses on pro-tumor functions for T cells in the SCLC environment. Such pro-tumor functions of T cells have been identified before, including in the context of chronic inflammation in colon cancer ^82^. as well as with immunosuppressive Tregs ^83^, a T-cell type that may contribute to the immune-cold environment in SCLC ^84,85^. Production of pro-inflammatory cytokines by activated T cells may also remodel the tumor environment and promote the expansion of cancer cells indirectly (e.g., IL-17 in a melanoma model ^86^); this could be important in SCLC, where chronic inflammation can be prevalent due to smoking or pollution ^87,88^. Our work identifies a new pro-tumor mechanism linked to T cells, via the production of IL-6, which serves as a pro-survival signal in SCLC cells. A screen of responses of human peripheral blood cells to a large set of cytokines did not identify a cytokine that can induce IL-6 expression in T cells ^89^. Future work will need to investigate how T cells in the SCLC microenvironment gain the ability to produce IL-6, and if these IL-6-producing T cells have additional properties relevant to tumor growth. In particular, future studies will need to investigate the interactions between T cells and other immune cells in the microenvironment of SCLC tumors, including γδ T cells, NK cells, and myeloid cells, which can play an important role in tumor growth and metastasis ^90,91^. Overall, tilting the balance towards favoring the anti-tumor effects of T cells may help enhance the survival of SCLC patients.

Our data show that CD74 expression is induced by TILs cells in SCLC. A recent study in colorectal cancer showed a similar induction of CD74 in cancer cells by T cells ^92^. Our work further identified CD74 as a key mediator of T cell-induced pro-tumor activity in SCLC. CD74 is primarily known as a key regulator of MHC II complex assembly and trafficking ^68^. However, as a receptor for MIF ^74^, CD74 can also serve as a pro-tumor receptor involved in anti-apoptotic signaling in cancer cell lines ^69–71^. Our data support a potent pro-survival role for MIF/CD74 signaling in SCLC. Notably, MIF/CD74 targeting is the focus of several pre-clinical and clinical approaches, in immune diseases and cancer ^93,94^. A previous study has suggested that inhibiting MIF/CD74 could enhance the effects of chemotherapy in SCLC cells in culture ^95^. Additionally, blocking CD74 might improve the anti-tumor effects of ICI in some contexts ^96^. Thus, targeting MIF/CD74 may be a promising strategy in SCLC. A CD74-blocking antibody has been tested in blood cancer and auto-immune disorders ^97,98^. However, because CD74 signaling plays a central role in antigen presentation and immune activation ^74,93,99^, it will be critical to further investigate the multiple effects that inhibition of MIF/CD74 signaling may have in SCLC before clinical strategies are implemented.

Our work points to IL-6 as a mediator of the pro-tumor functions of T cells in SCLC cells. Emerging evidence from SCLC patients supports this finding, in addition to our analysis of the IMpower133 clinical trial data. For instance, high levels of IL-6 in the blood of SCLC patients has been associated with resistance to ipilimumab (targeting CTLA-4) ^100^. In addition, under current standard chemotherapy, SCLC patients with higher IL-6 expression level have a shortened overall survival ^101^. Similarly, SCLC patients with lower levels of IL-6 (as well as TNF-α and IFN-γ) show longer progression-free survival ^88^. A major difference between our observations and previous observations linking IL-6 blockade to improve responses to immunotherapies ^77–79^ is that, in our studies, IL-6 blockade directly affects the survival of cancer cells rather than immune cells in the tumor microenvironment. Together, these observations highlight the need to further investigate IL-6 in SCLC, and whether it comes from T cells or other cells in tumors ^60,61^. IL-6 is a particularly interesting target because there are FDA-approved therapeutics to block IL-6 signaling in patients, including via anti-IL-6 receptor monoclonal antibodies (e.g., tocilizumab, satralizumab) or anti-IL-6 monoclonal antibodies (i.e., siltuximab, ziltivekimab) ^102^. It is also noteworthy that blocking IL-6 signaling is already used as a strategy in the clinic to limit the cytokine release storm that can result from tumor killing by CAR T cells ^103^; as more CAR T cells and bispecific T cell engagers are being developed in SCLC ^81^, IL-6 pathway inhibitors may naturally be tested in the near future in this patient population. More broadly, recent evidence indicates that blocking signaling downstream of cytokines may enhance the efficacy of ICI in other cancer types ^104^. Ultimately, adding IL-6 pathway inhibitors to standard-of-care therapy, including ICI, may help extend the survival of patients with SCLC.

One experimental challenge with IL-6 is that mouse IL-6 is not active on human cells ^105^, which limits studies of human tumors growing in mice and highlights the need of relevant pre-clinical models. One additional limitation of our work is that we only used IL-6 blockade in our pre-clinical models: it is possible that strategies targeting the IL-6 receptor would have greater therapeutic effects, although it is also possible that side effects may be greater. Indeed, in the clinic, targeting IL-6 has been associated with a greater risk of serious infections ^102^; this may not be a critical issue in patients with SCLC because these patients undergoing chemotherapy are already under surveillance for these side effects, but this remains a possible limitation.

## Methods

### Ethics statement

Mice were maintained according to practices approved by the NIH, the Stanford Institutional Animal Care and Use Committee (IACUC), and the Association for Assessment and Accreditation of Laboratory Animal Care (AAALAC). The study protocol was approved by the Stanford Administrative Panel on Laboratory Animal Care (APLAC) (protocol 13565).

### Animal studies

For tumor induction in the *RPR2* model (*RPR2-Luc*), 8-12 weeks old male and female mice in a mixed 129Sv/J and C57BL/6 background were infected by intratracheal instillation with 2×10^10^ PFU/mL of Adeno-CMV-Cre (Baylor College of Medicine). B6129SF1/J mice (Jackson Laboratories, stock no. 101043) were used for allograft experiments in immunocompetent recipients. Nod.Cg-Prkdc^scid^IL2rg^tm1WjI^/SzJ (NSG) mice were used for experiments in immunodeficient mice (Jackson Laboratories, stock no. 005557). Mice were engrafted with 0.25-1×10^6^ cancer cells in antibiotic-free serum-free medium with a 1:1 mixture of Matrigel (BD Matrigel, 356237) at 8-12 weeks of age. Both male and female mice were used. Therapeutic agents were administered by intraperitoneal injection. Tumor volumes were calculated as 0.5 × length × width^2^. Mice were housed at 22°C ambient temperature with 40% humidity and light/dark cycle of 12h (7 a.m.-7 p.m.).

For T cell depletion experiments in the *RPR2* model, mice were imaged 4 months post intratracheal injection for luciferase signal using an SII Lago-X instrument. Once the presence of tumors was confirmed, mice were randomized into two groups (based on mean tumor growth) and treated with PBS or 20 µg anti-mouse CD3ε antibody (BE0001-1, Bio X Cell), 3 times a week for one month. For allograft models, tumors were allowed to grow for 7-10 days and then mice were randomized into treatment groups with PBS or 100 µg anti-mouse CD3ε antibody 3 times a week, for 2-3 weeks. Tumor size was measured 2 times a week.

For T cell transfer experiments, T cells were isolated using the EasySep™ Mouse T Cell Isolation Kit (19851, STEMCELL) from the lungs and spleen of *RPR2* mutant mice ∼6-7 months after tumor induction. Lungs were finely chopped with a razor blade and digested in 25 mL of L-15 Medium (Leibovitz) (L1518, Millipore Sigma) media with a five-enzyme mix containing DNase I (10104159001, Roche), Elastase (LS002279, Worthington Biochemical Company), Collagenase Type I (C0130, Millipore Sigma), Collagenase Type II (C6885, Millipore Sigma), Collagenase Type IV (C5138, Millipore Sigma) for 45 min on a shaker at 37°C. The digested mixture was then passed through a 40 µm filter, spun down, washed with 10 mL PBS, resuspended with 5 mL of red blood lysis buffer (00-4333-57, eBioscience™ 1X RBC Lysis Buffer) for 1 min at room temperature and stopped by adding 35 mL PBS, and washed again with 10 mL PBS. Spleens were manually crushed and then filtered, washed, lysed for red blood cells, and washed again in the same manner as the lungs. T cells were isolated from both lungs and spleen single cells using the EasySep™ Mouse T Cell Isolation Kit (19851, STEMCELL). SCLC cells were injected alone or with T cells from the spleen or the lungs (ratio of 1 T cells for 10 SCLC cells) into the flanks of NSG mice.

For the combination treatment experiments, mice were randomized into treatment groups: PBS, 100 µg anti-IL-6 (BE0046, Bio X Cell), or 100 µg anti-PD-1 (BE0146, Bio X Cell), which were injected 3 times a week for 2-3 weeks.

### Cell lines

*Rb/p53* mutant mouse SCLC KP1 cells and *Rb1/Trp53/Rbl2* mutant mouse SCLC KP11 cells were described previously; both cell lines express ASCL1, and thus represent the most common subtype of SCLC ^51^. The genetic background of these mouse cell lines is a mix between C57BL/6 and S129/Sv. All SCLC cells were cultured in RPMI-1640 supplemented with 10% fetal bovine serum (Hyclone), 1× GlutaMax (Invitrogen) and 100 U/mL penicillin and 100 µg/mL streptomycin (Invitrogen). Cell lines were grown in suspension and dissociated by gently pipetting. Cell lines were cultured in humidified incubators at 37°C with 5% carbon dioxide. All cell lines were routinely tested (Lonza) and confirmed to be free of mycoplasma contamination.

### Cytokine profiling

For cytokine profiling 25,000 T cells isolated from the lung and spleen of SCLC tumor-bearing mice were incubated for two days in 200 µL of RPMI medium containing fetal bovine serum, glutamine, penicillin, and streptomycin. T cell conditioned media were then stored at -80°C before analysis using the mouse Luminex 48-plex cytokine array (Procarta) at the Stanford University Human Immune Monitoring Center.

### Knock-down and Knock-out

CD74 control and knock-down cells were generated using a control plasmid (Addgene, 8453) and two predesigned shRNA plasmids (Millipore Sigma, TRCN0000066091 and TRCN0000066092). Control and *gp130* knock-out cells were generated using pre-purchased Cas9 and single-guide plasmids (VectorBuilder, VB010000-9355sqw, VB900129-9335mkp, VB900129-9336bpj). Control and *Il-6st* knock-down cells were generated using two predesigned shRNA plasmids (Millipore Sigma, TRCN0000065391 and TRCN0000348799). To generate knock-down or knock-out cells, lentiviral supernatants were prepared by transfecting 15×10^6^ HEK293T cells seeded in 15-cm plates and transfected with a mixture of the lentivirus vector plasmids (1:1:1 of VSV-G, pMDLg/pRRE, and pRSV/Rev) with Lipofectamine™ 3000 (Thermo Fisher Scientific, L3000075). The virus was then filtered for residual HEK293T cells and was incubated with KP1 or KP11 cells for 24 hr with 8 µg/mL of polybrene. After 24 hr, cells were incubated with fresh medium containing 1 µg/mL puromycin (Thermo Fisher Scientific, A1113803) for 3 days. Selected cells were recovered in fresh media for 2 days before analysis.

### Immunostaining

For immunofluorescence staining on paraffin sections, mouse tissues were first fixed overnight using 4% paraformaldehyde and processed for paraffin embedding. Paraffin sections were deparaffinized with Histo-Clear (National Diagnostics HS-200) and rehydrated. Antigen retrieval was carried out in a citrate-based unmasking solution (Vector Laboratories H-3300) by heating in the microwave till boiling followed by an additional 12 min at 30% power. Sections were washed in PBST (PBS with 0.1% Tween-20) and blocked in 5% horse serum for 1 hr. Primary antibodies diluted in PBST with 5% horse serum were then added to the sections for incubation overnight at 4°C. After washing, a secondary fluorescent antibody was applied and incubated for 1 hr at room temperature. Nuclei were counterstained with DAPI (D9542, Sigma-Aldrich), and slides were mounted with ProLong™ Diamond Antifade Mountant (P36961, Thermo Fisher Scientific). Images were captured using Keyence BZ-X700 all-in-one fluorescence microscope with BZX Viewer program version 1.3.1.1. Tumor area, CD74, and DAPI staining were quantified using the ImageJ 1.52K software. The following antibodies were used for mouse sections: anti-CD74 (ab245692, Abcam), anti-mouse CD3 (14-0032-85, Thermo Fisher Scientific), anti-mouse CD4 (MA1-146, ThermoFisher), anti-mouse CD8 (MA1-145, Thermo Fisher Scientific), anti-IL-6 (P620, ThermoFisher), Alexa Fluor™ 594 Goat anti rabbit (A-11037, Thermo Fisher Scientific), Alexa Fluor® 488 donkey anti-rat (A-21208, Thermo Fisher Scientific).

For human SCLC tumor analysis, tissue microarrays (LC802 and LC245, Tissue Array). Staining was performed in a similar manner as written above. The following antibodies were used: anti-CD74 (ab245692, Abcam), anti-human CD3 (MB600-1441, Novus Biologicals), anti-IL-6 (P620, ThermoFisher), Alexa Fluor™ 594 Goat anti rabbit (A-11037, ThermoFisher), Alexa Fluor® 488 donkey anti-rat (A-21208, ThermoFisher).

### Flow cytometry

For cell surface flow cytometry analysis, single-cell suspensions were resuspended in PBS and counted. Fc receptors were blocked with anti-mouse CD16/32 antibody (101302, BioLegend), followed by staining with a conjugated antibody cocktail for 30 minutes at 4°C. The following antibodies were used: FITC rat IgG1 Isotype control Antibody (401913, BioLegend), APC rat IgG1 Isotype control Antibody (401903, BioLegend), PE anti-mouse CD130 (149403, BioLegend), Pacific Blue anti-mouse CD3 (100214, BioLegend), FITC anti-mouse CD4 (100406, BioLegend), Brilliant Violet 605 anti-mouse CD4 (100451, BioLegend), PE-Cy7 anti-mouse CD8α (100722, BioLegend), PE anti-mouse CD8α (100707, BioLegend), Brilliant Violet 650 anti-mouse CD279 (135243, BioLegend), APC anti-mouse CD39 (143809, BioLegend), PE/Cyanine7 anti-mouse/human CD11b (101215, BioLegend), Brilliant Violet 421 anti-mouse/human CD11b (101235, BioLegend), PE/Cyanine7 anti-mouse CD62L (304821, BioLegend), Brilliant Violet 510 anti-mouse/human CD44 (103043, BioLegend), and Zombie NIR Fixable Viability Kit (423105, BioLegend) or DAPI (D9542, Sigma-Aldrich) for live/dead discrimination.

For Intracellular IL-6 Staining, cells were treated with a Cell Stimulation Cocktail plus protein transport inhibitors (00-4975-93, Invitrogen) for 2 hours at 37°C. Cells were then first stained for surface markers as described above, followed by incubation with the Zombie NIR™ Fixable Viability Kit (423105, BioLegend) to exclude dead cells. Cells were subsequently fixed and permeabilized using the BD Cytofix/Cytoperm™ Kit (554714, BD Biosciences) according to manufacturer’s instructions. Intracellular staining was performed overnight at 4°C using anti-mouse IL-6 antibody (561367, BD Biosciences) diluted in BD Perm/Wash™ buffer.

After staining, cells were washed and resuspended for flow cytometry analysis. Data were acquired using either a BD LSRFortessa™ Cell Analyzer or BD FACSDiscover™ S8 Cell Sorter. Flow cytometry data were analyzed and visualized using FCS Express software.

### Cell cycle and cell death analyses

Cell death was measured by Annexin V and propidium iodide (PI) staining. Accutase (423201, BioLegend) treatment for 1 min was used to generate single-cell suspensions before a PBS wash and resuspension in the Annexin V buffer (422201, BioLegend). The cell pellets were resuspended with APC-Annexin V (550474, BD Biosciences). Cells were incubated for 30 min in the dark on ice before washing and analysis. For cell cycle analysis, cells were fixed using 1.6% PFA for 15 min on ice and washed twice with PBS before staining with the FxCycle™ PI/RNase Staining Solution (F10797, ThermoFisher) and analysis.

### Immunoassays and protein extraction

Cells were lysed in RIPA buffer (89900, Thermo Fisher Scientific) supplemented with protease and phosphatase inhibitors (05892970001, Roche cOmplete ULTRA tablets, Mini, EASYpack and 04906845001, PhosSTOP EASYpack). Protein concentration was measured using a PierceTM BCA protein assay kit (23227, Thermo Fisher Scientific). Immunoassays were conducted using the capillary-based Simple Western^TM^ assay on the Wes^TM^ system (ProteinSimple). Compass software v4.0.0 was used to analyze and visualize the results. The following antibodies were used: HSP90 (4877S, Cell Signaling), CD74 (AF7478, R&D Systems), STAT3 (4904, Cell Signaling), phospho-STAT3 (9145, Cell Signaling), Akt (4691, Cell Signaling), Phospho-Akt (4058, Cell Signaling).

### Cell viability assay

Cell viability was measured using the alamarBlue™ Reagent (ThermoFisher, DAL1100). 5,000 KP1, KP11, NCI-H69 or NCI-H82 were seeded in 100 µL of medium. ISO-1 (Selleckchem, S7732) recombinant mouse IL-6 (R&D systems, 406-ML) were added as needed. Signal intensity was measured using the Synergy H1 Hybrid Redear (BioTekTM) after adding 10 µL of alamarBlue reagent to each well and incubating for 4 hr in 37°C.

### Bulk RNA sequencing

Digested subcutaneous tumors from mice were filtered (40 µm filter), spun down, washed with 10 mL PBS, and resuspended with 5 mL of red blood lysis buffer (00-4333-57, eBioscience™ 1X RBC Lysis Buffer) for 1 minute at room temperature and stopped by adding 35 mL PBS. Pellets of 10^6^ cells were flash frozen after one more wash before RNA sequencing. Cells in culture were spun down and flash-frozen before RNA sequencing. Library preparation and sequencing were done by Novogene. RNA counts were quantified using Salmon and differential RNA-seq analysis was conducted using DESeq2 as previously described ^106^.

### Single-cell RNA sequencing

T cells characterization in *RPR2* mutant mouse model: T cells were isolated from the lungs and spleen of *RPR2* mutant mice (n=3) using flow cytometry sorting with a CD3 antibody. 10,000 cells per sample were barcoded and libraries were generated using the V2 10x Chromium system. The samples were sequenced using NovaSeq with a target of 30,000 reads per cell. Before analysis, data from each sample were individually pre-processed using the CellRanger pipeline as a Seurat object ^107^. Then for each Seurat object, low quality cells, doublets, cells with low mitochondrial reads (< 15) and cells with <250 & > 6000 expressed genes were removed. After removing unwanted cells from the dataset, normalization of the data was performed using the SCTransform function ^108^. Lastly, the two spleen and lung Seurat objects were integrated using the FindIntegrationAnchors function of Seurat.

Cluster identity was determined by the following markers ^109^: main CD8 T cells expressing *Cd8a*, CD4 T cells expressing *Cd4*, naive/central memory cells expressing *Tcf7* and *Ccr7* while lacking activation features such as *Pdcd1*; effector memory T cells expressing *Tcf7* and granzymes (*Gzmb* and *Gzmk*); resident memory T cells (Trm) expressing *Cxcr6* and *Itga1* and low levels of granzymes ^110,111^; effector T cells expressing granzymes and perforins (*Prf1)* including a unique subpopulation of cytotoxic CD8 T cells expressing *Cx3cr1* ^112^; exhausted (Tex) effector cluster with high expression of granzymes and inhibitory receptors including *Ctla4*, *Lag3*, and *Entpd1*; regulatory T cells (Treg), identified by *Foxp3;* Th2 CD4 T cells expressing *Gata3* and *Il4* ^113^; double-negative T cells identified by low expression of *Cd8a* and *Cd4* expression; gamma delta T cells were identified by expression of *Trdc*, and alpha-beta T cells by *Trac*.

IL-6 and PD-1 blockade combination treatment: single-cell analysis of KP11 allograft tumors treated with IL-6 blockade alone, PD-1 blockade alone, or combination treatment of anti-PD1 and IL-6 blockade was done as described above. At the experimental endpoint, tumors were harvested and dissociated. Due to the low representation of immune cells in KP11 allografts, CD45⁺ cells were enriched using the EasySep™ Mouse CD45 Positive Selection Kit (18945, STEMCELL Technologies).

For single-cell library preparation, cells were fixed and barcoded using Parse Biosciences Evercode Fixation and Evercode WT kits, respectively, according to the manufacturer’s instructions. Sequencing was performed on a NovaSeq.

Before analysis, data from each sample were individually pre-processed in a similar manner as described before. For cell-type identification, immune cells were defined as CD45⁺ cells, fibroblasts were defined by *Pdgfra* expression, endothelial cells were defined by expression of *Pecam1*, *Cdh5*, *Cldn5*, and *Erg*, and SCLC cells were defined by negative selection (*CD45⁻ Pdgfra⁻ Pecam1⁻ Mrc1⁻ Itgam⁻ Adgre1⁻ Cd163⁻*). For immune cell subpopulation identification, the following markers were used: T cells were identified by expression of *Cd3e*, *Cd3g*, *Cd4*, and *Cd8a*. Macrophages were identified by expression of *Adgre1*, *Itgam*, *Cd68*, *Mrc1* and *Cd163*. Dendritic cells were identified by expression of *Itgax*, *H2-Ab1*, and *Cd74*. Natural killer cells were identified by expression of *Ncr1*, *Klrb1c*, and *Klrc1*. B cells were identified by expression of *Cd19*. Granulocytes were identified by expression of *Ly6g*, *S100a8*, and *S100a9*. For T cell functional analysis, cytotoxic markers were defined by expression of *Prf1*, *Gzmb*, *Gzmk*, and *Gzma*, while exhaustion markers were defined by expression of *Pdcd1*, *Lag3*, *Ctla4*, and *Entpd1*.

For SCLC cell analysis, the inflammatory response was assessed using Hallmark gene sets. MHC-I expression was evaluated using the signature genes *H2-K1*, *H2-D1*, *B2m*, *Tap1*, *Tap2*, *Psmb8*, *Psmb9*, and *Tapbp*. AKT pathway activity was defined by expression of *Pik3ca*, *Akt1*, *Akt2*, *Akt3*, *Pdk1*, *Pten*, and *Mtor*. FAS-mediated apoptosis was assessed using the REACTOME_EXTRINSIC_PATHWAY_FOR_APOPTOSIS gene set, including *Fas*, *Faslg*, *Fadd*, *Casp8*, *Casp10*, *Casp3*, *Casp7*, *Cflar*, *Bid*, and *Bax*. NF-κB pathway activity was evaluated using genes *Traf1*, *Traf2*, *Rela*, *Birc3*, *Relb*, *Nfkb2*, and *Cd40* ^72,114^. Neuroendocrine (NE) and non-neuroendocrine genes were defined as described previously ^27^.

### Analysis of IMpower133 RNA-seq data

Previously published bulk RNA-seq data from IMpower133 trial were used for analyses ^26^. Median gene signature expression was used to subset patients into gene-high v. gene-low cohorts. For signature score determination, expression levels of genes in the signature were first converted to Z-scores using the median, and the mean Z-score of the signature genes was used to determine the overall signature Z-score. Median signature Z-score was similarly used to separate signature-high and signature-low cohorts. A previously published T cell signature (*TBX21*, *ITK*, *CD3D*, *CD3E*, *CD3G*, *TRAC*, *TRBC1*, *TRBC2*, *CD28*, *CD5*, and *TRAT1*) was used ^115^. Cox proportional hazard ratios and forest plots were calculated and generated using R (4.3.1) package survival (3.5-8) and forestplot (3.1.3), respectively.

### Statistical and reproducibility

Statistical significance was assayed with GraphPad Prism software version 10.2.3. Data are represented as mean ± s.e.m. Tests used are indicated in figure legends. Data distribution was assumed to be normal, but this was not formally tested. No data were excluded from the analysis. To compare two groups, we used a t-test. To compare growth curves, we used two-way ANOVA.

## Data availability

Sequencing data from RNA sequencing are available from the Gene Expression Omnibus (GEO) under accession numbers GSE290290. All other data are available in the article and supplementary materials, or from the corresponding author upon reasonable request.

## Acknowledgements

We thank all the members of the Sage lab for their help and support throughout this study, Dr. Alison Farrell for her critical reading of the manuscript, and Drs. Alex Jaeger, Nikhil Joshi, and Peter Westcott for helpful discussions. Research reported in this publication was supported by the NIH (J.S. R35 CA231997), a Mark Foundation Endeavor Award, and the California TRDRP (M.B. T32FT4934), J.S. is the Elaine and John Chambers Professor in Pediatric Cancer.

---

**Supplementary Figure S1, related to Figure 1:**
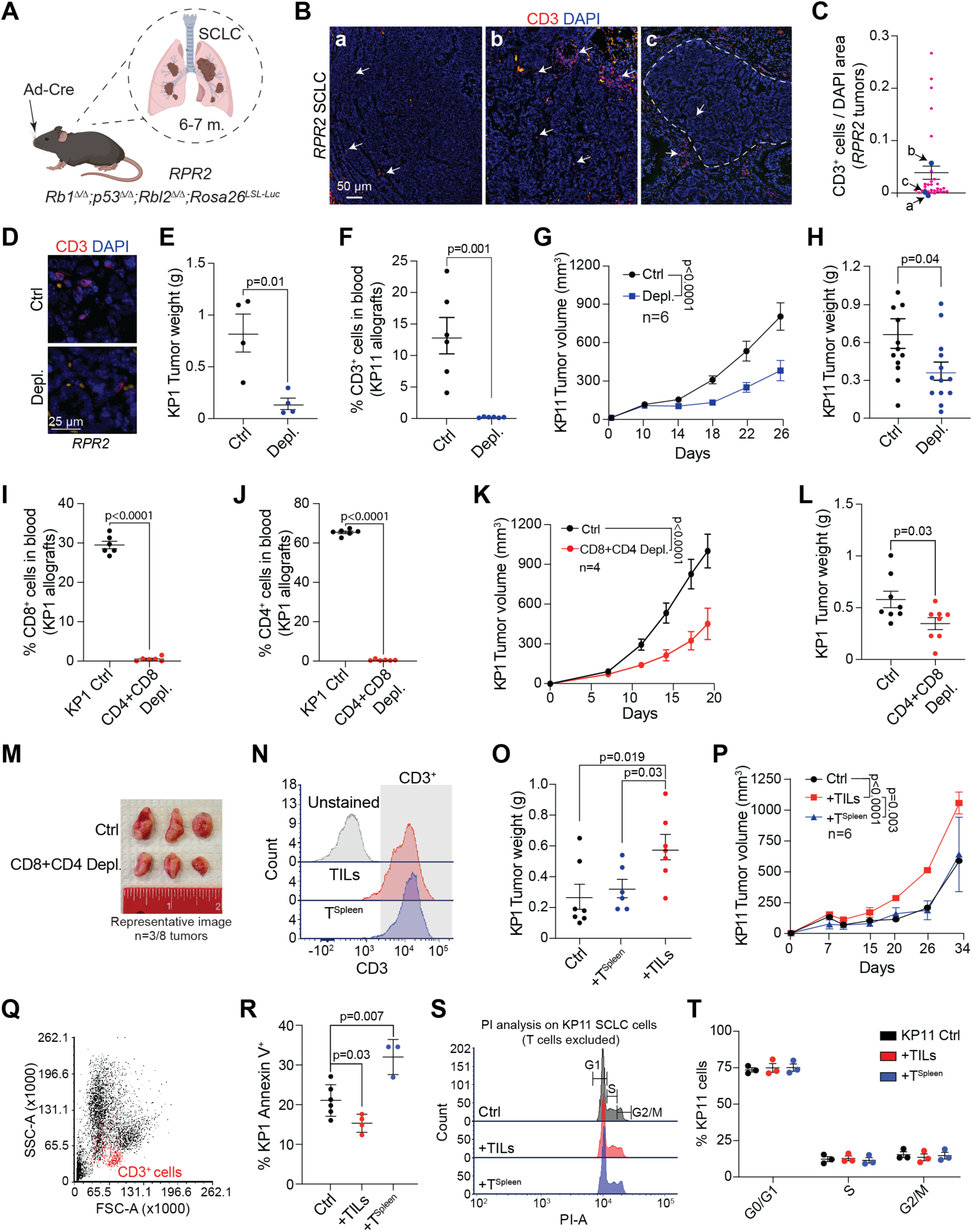
T cells from the SCLC microenvironment promote tumor growth. **A.** Schematic illustrating the *RPR2* autochthonous mouse model of SCLC. **B.** Representative immunofluorescence staining of CD3^+^ T cells (red) in *RPR2* tumors. Nuclei are counterstained with DAPI (blue). Scale bar, 50 µm. White arrows indicate CD3^+^ T cells. Examples of SCLC tumors with low (a), medium-high (b) T cell infiltration, and peritumoral T cells (c) are shown. **C.** Quantification of T cells from SCLC tumors in the *RPR2* model, calculated as the ratio of CD3^+^ signal to DAPI signal in tumor sections, as shown in (B) (n=9 mice). Data for (a), (b), and (c) correspond to cases in (B). Error bars represent s.e.m. **D.** Representative immunofluorescence staining of CD3^+^ T cells (red) in SCLC tumors from control (Ctrl) and T cell-depleted (Depl.) *RPR2* mice (n=5). Nuclei are counterstained with DAPI (blue). Scale bar, 25 µm. **E.** Tumor weight for KP1 SCLC allografts in control (Ctrl) and T cell-depleted (Depl.) mice (n=4 mice). Statistical significance was determined by a t-test; error bars represent s.e.m. **F.** Percentage of CD3^+^ T cells in the peripheral blood of control (Ctrl) and T cell-depleted (Depl.) mice bearing KP11 SCLC allografts (n=6). Statistical significance was assessed using a t-test; error bars represent s.e.m. **G.** Tumor growth curves of KP11 SCLC allografts in control (Ctrl) and T cell-depleted (Depl.) mice (n=4, n=2 tumors per mouse, n=1 experiment). Statistical analysis was performed using two-way ANOVA; error bars represent s.e.m. **H.** Tumor weight in grams for KP11 allografts from control (Ctrl) and T cell-depleted (Depl.) mice as in (G). Statistical significance was assessed with a t-test; error bars represent s.e.m. **I, J.** Percentage of CD8^+^ (I) and CD4^+^ (J) T cells in the blood of control (Ctrl) and CD4+CD8-depleted (Depl.) mice bearing KP1 SCLC allografts (n=4, 2 tumors per mouse, n=1 experiment). Statistical significance was assessed using a t-test; error bars represent s.e.m. **K.** Tumor growth curves of KP1 SCLC allografts in control (Ctrl) mice and mice with depletion of both CD4^+^ and CD8^+^ T cells (CD4+CD8 Depl.) (n=4, 2 tumors per mouse, n=1 experiment). Statistical analysis was performed using two-way ANOVA; error bars represent s.e.m. **L.** Tumor weight at terminal time point for KP1 SCLC allografts as in (K). Statistical significance was assessed by a t-test; error bars represent s.e.m. **M.** Representative images of KP1 SCLC tumors at the terminal time point from control (Ctrl) and CD4+CD8-depleted (CD4+CD8 Depl.) mice, as in (K). **N.** Representative flow cytometry analysis showing CD3^+^ cell purity following isolation of TILs and splenic T cells (T^Spleen^) from tumor-bearing *RPR2* mice, as in Fig. 1I-K. **O.** Tumor weight for KP1 allografts either injected alone (Ctrl) or co-injected with T cells isolated from spleen (T^Spleen^) or SCLC tumors (TILs) of tumor-bearing *RPR2* mice as in Fig. 1I-K. Statistical significance: a t-test; error bars, s.e.m. **P.** Tumor growth curves of KP11 SCLC allografts in NSG mice injected alone (Ctrl) or with T cells isolated from the spleen (T^Spleen^) or SCLC tumors (TILs) of tumor-bearing *RPR2* mutant mice (n=6 mice, n=1 tumor per mouse, n=1 experiment). Statistical analysis was performed using two-way ANOVA; error bars represent s.e.m. **Q.** Representative flow cytometry plot (SSC-A vs. FSC-A) of SCLC cell line co-cultured with T cells (red). **R.** Apoptosis analysis of KP1 SCLC cells cultured alone (Ctrl) or co-cultured with tumor-infiltrating lymphocytes (TILs) or splenic T cells (T^Spleen^) for 48 hours. Apoptosis was analyzed only in CD3-negative cells. Statistical significance was assessed using a t-test; error bars represent standard deviation (n≥3 independent experiments). **S.** Representative flow cytometry analysis (propidium iodide, PI) of KP11 cells grown alone or with T cells isolated from the spleen (T^Spleen^) or SCLC tumors (TILs). **T.** Quantification of (S) (n=3). Statistical significance was assessed using a t-test (not significant); error bars represent s.e.m.

**Supplementary Figure S2, related to Figure 2:**
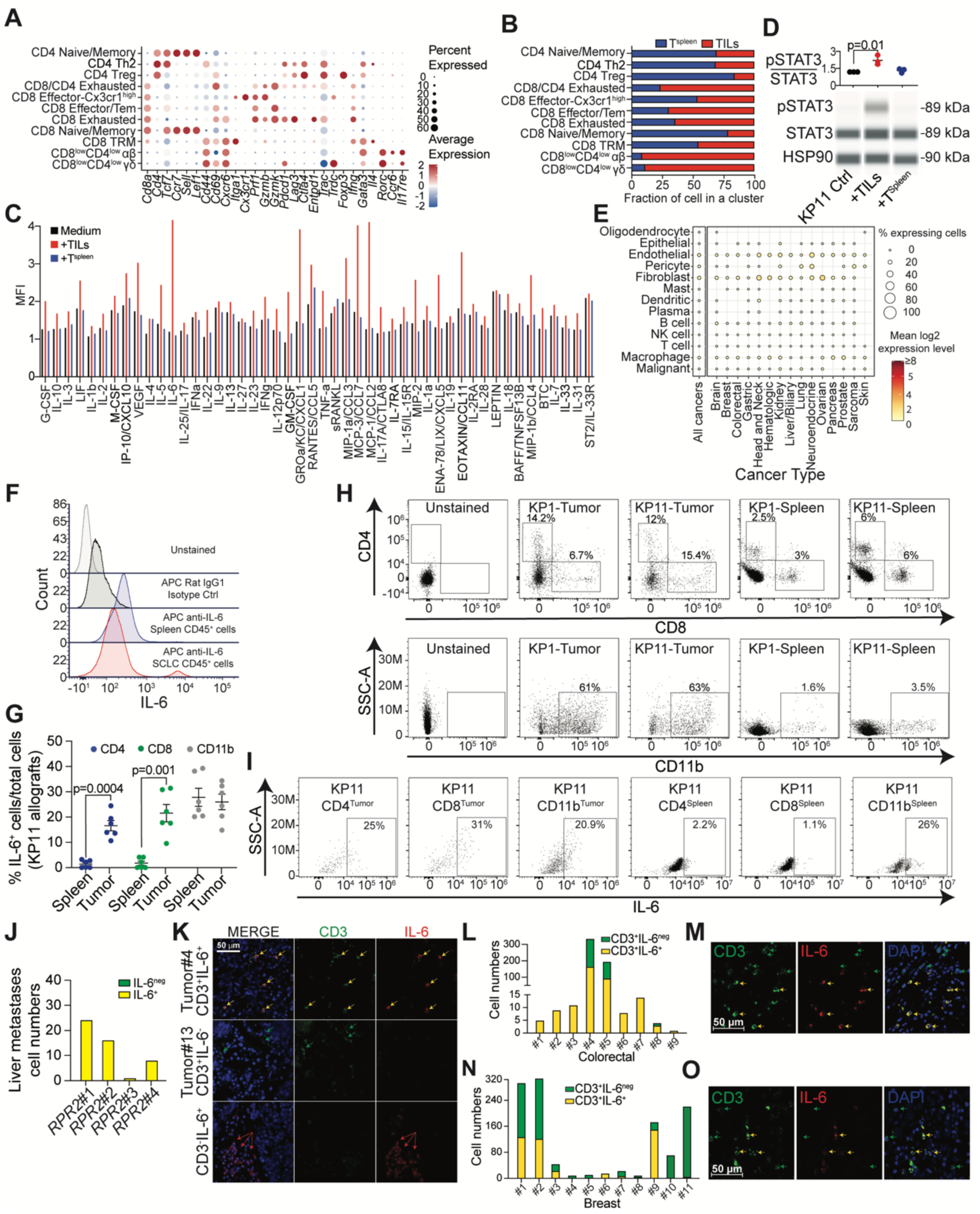
A population of T cells express IL-6 in SCLC tumors. **A.** Dot plot of representative marker genes for T cell populations identified by scRNA-seq in Fig. 2A. Circle size reflects the percentage of cells expressing each gene, and color intensity represents average expression level. **B.** Fraction of splenic T cells (T^Spleen^) or TILs from each cluster as in (A). (n=3). **C.** Mean Fluorescence Intensity (MFI) for 48 cytokine levels in the supernatant of T^Spleen^ cells and TILs cells isolated from tumor-bearing *RPR2* mutant mice and kept in culture for 48 hours. **D.** Representative immunoblot and quantification of STAT3 and phosphorylated STAT3 (pSTAT3) levels normalized to HSP90 in KP1 SCLC cells cultured alone or co-cultured with TILs or splenic T cells (T^Spleen^). Statistical significance was assessed by a t-test; error bars represent SD (n=3). **E.** Pan-cancer single-cell RNA-seq analysis for *IL-6* expression from ^116^. **F.** Representative flow cytometry plots showing intracellular IL-6 staining in CD45^+^ cells isolated from the spleen or SCLC tumors of *RPR2* mutant mice. Negative controls include unstained (no antibody) and rat IgG1 isotype control samples. **G.** Percentage of IL-6-expressing cells within total CD8⁺ (green), CD4⁺ (blue), and CD11b⁺ (gray) populations in spleen versus KP11 SCLC allograft tumors from tumor-bearing mice (n=5 mice). Statistical significance was determined by a t-test; error bars represent s.e.m. **H.** Representative flow cytometry demonstrating gating strategy (top row) for CD4⁺ and CD8⁺ T cells isolated from KP1 and KP11 SCLC allograft tumors (Tumor) and splenic cells (Spleen). CD11b⁺ populations (bottom row) in KP1 and KP11 SCLC allograft tumors (Tumor) and spleen (Spleen). **I.** Representative flow cytometry analysis of IL-6 expression in unstained controls, CD4⁺ cells, CD8⁺ cells, and CD11b⁺ cells isolated from SCLC tumors (Tumor) or splenic cells (Spleen) of KP11 allograft-bearing mice. **J.** Quantification of IL-6^+^ and IL-6^neg^ CD3^+^ T cells in liver metastases of *RPR2* mutant mice (n=4). **K.** Representative immunofluorescence images of human SCLC tumor sections. IL-6 is shown in red; CD3 in green; nuclei are counterstained with DAPI (blue). Yellow arrows indicate IL-6⁺CD3⁺ double-positive cells (tumor #4); green arrows indicate CD3⁺ single-positive cells (tumor #13); red arrows indicate CD3^neg^IL6^+^ cells. Scale bar, 50 µm. **L.** Quantification of IL-6⁺ and IL-6^neg^ CD3⁺ T cells in human colorectal tumor sections by immunofluorescence. **M.** Representative immunofluorescence images of human colorectal tumor sections. IL-6 is shown in red; CD3 in green; nuclei are counterstained with DAPI (blue). Yellow arrows indicate IL-6⁺CD3⁺ cells; green arrows indicate CD3⁺ single-positive cells. Scale bar, 50 µm. **N.** Quantification of IL-6⁺ and IL-6^neg^ CD3⁺ T cells in human breast tumor sections by immunofluorescence. **O.** Representative immunofluorescence images of human breast tumor sections. IL-6 is shown in red; CD3 in green; nuclei are counterstained with DAPI (blue). Yellow arrows indicate IL-6⁺CD3⁺ cells; green arrows indicate CD3⁺ single-positive cells. Scale bar, 50 µm.

**Supplementary Figure S3, related to Figure 3:**
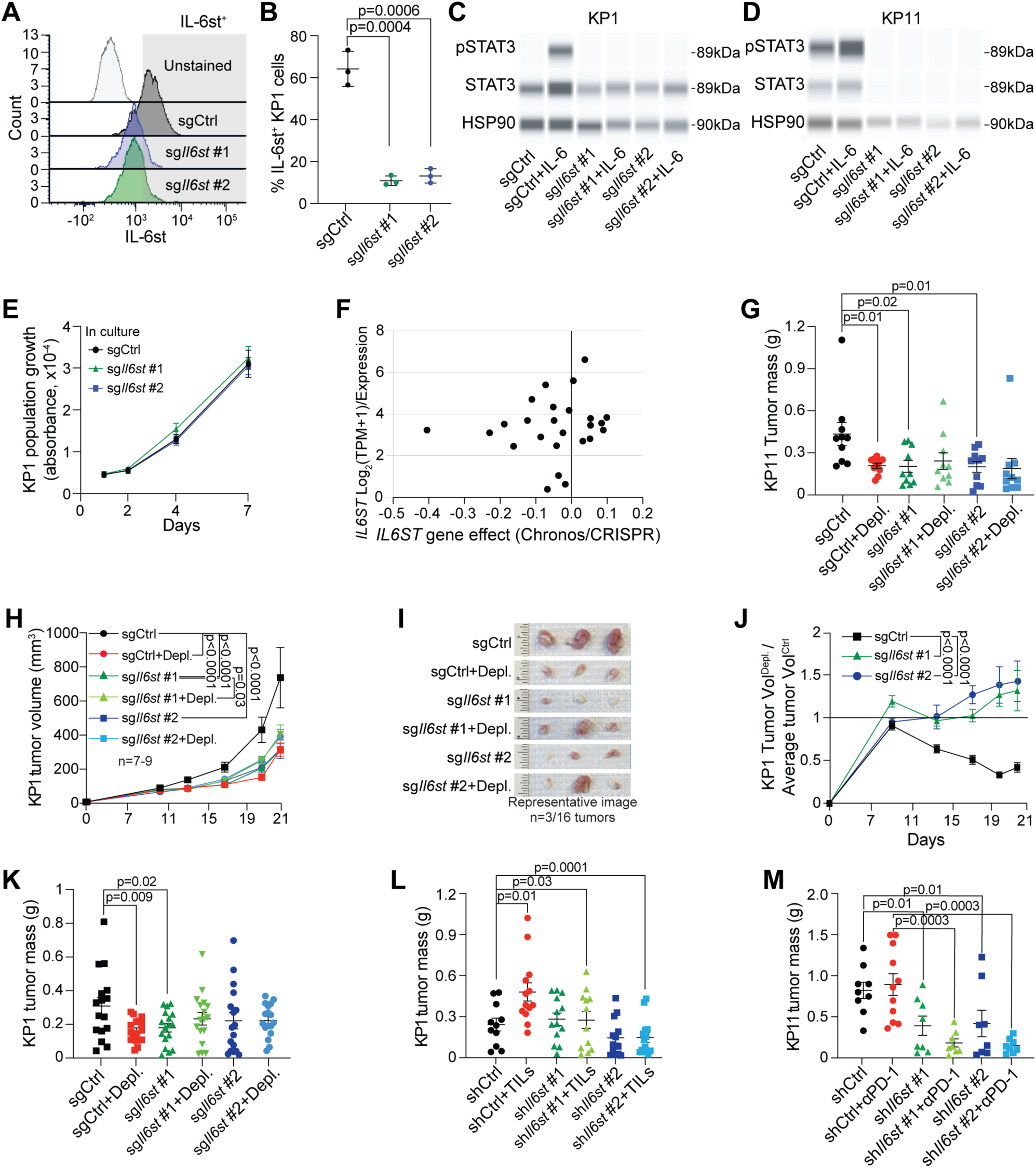
T cells promote tumor growth via the IL-6 receptor in SCLC cells. **A.** Representative flow cytometry analysis for IL-6st levels in control (sgCtrl) or *Il6st* knock-out KP11 cells (sg*Il6st* #1 and #2). Unstained cells provide a negative control for gating. **B.** Quantification of IL-6st expression from (A) (n=3). Statistical significance was determined by t-test; error bars represent SD. **C, D.** Immunoassay for STAT3 and phosphorylated STAT3 (pSTAT3) levels relative to HSP90 in KP1 (C) and KP11 (D) mouse SCLC allografts with control of *Il6st* knock-out, in the presence or absence of recombinant IL-6. **E.** Cell viability assay (AlamarBlue) of control (sgCtrl) and *Il6s*t knock-out (sg*IL6st* #1 and #2) KP11 cells for 7 days (n=3). Statistical analysis was performed using two-way ANOVA; no significant differences were observed. Error bars represent SD. **F.** DepMap data representing the levels of expression of *ILST6* as a function of the effects of its knock-out in a collection of human SCLC cells. Positive gene effect values indicate a pro-growth phenotype while negative gene effect values indicate an anti-growth phenotype. **G.** Tumor weight for KP11 SCLC allografts as in Fig. 3D (terminal time point) (n=10). Statistical significance was assessed using a t-test; error bars represent s.e.m. **H.** Tumor growth curves of KP1 SCLC tumors, with or without *Il6st* knock-out, in control and T cell-depleted (+Depl.) mice (n=7-9 mice, n=1-2 tumors per mouse, n=1 experiment). Statistical analysis was performed using two-way ANOVA; error bars represent s.e.m. **I.** Representative images of KP1 tumors at the terminal time point, as in (H). **J.** Fold change in tumor size from (H). Tumor volume at each time point was normalized to the mean of control tumors. Ratio >1 indicates an anti-tumor effect of T cells, ratio =1 indicates no effect, and ratio <1 indicates a pro-tumor effect. As expected, control tumors showed ratio <1 (T cell depletion slowed growth), while knock-out tumors showed ratios closer to or >1. Statistical analysis by two-way ANOVA; error bars represent s.e.m. **K.** Tumor weight of KP1 SCLC allografts at endpoint, as in (H). Statistical significance by t-test; error bars represent s.e.m. **L.** Tumor weight for KP1 SCLC allografts, as in Fig. 3I (terminal time point) (n=7-8, n=1-2 tumors per mouse). Statistical significance was assessed using a t-test; error bars represent s.e.m. **M.** Tumor weight for KP11 SCLC allografts, as in Fig. 3L (terminal time point) (n=8-9, n=1-2 tumors per mouse). Statistical significance was assessed using a t-test; error bars represent s.e.m.

**Supplementary Figure S4, related to Figure 4:**
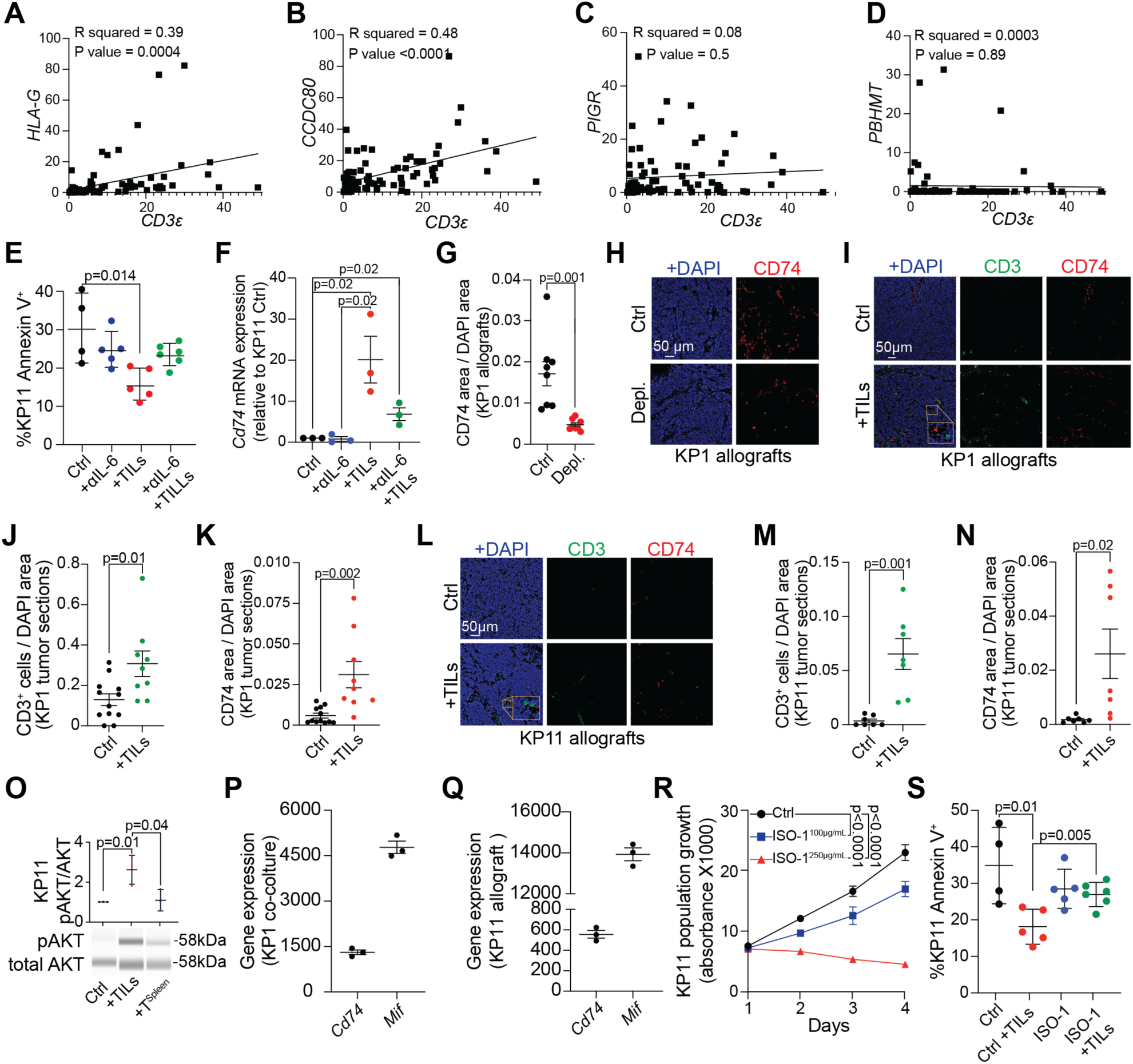
Expression of the pro-survival factor CD74 is induced in SCLC cells by TILs. **A-D.** Correlation of *CD3ε* (T cells) with expression of *HLA-G* (A), *CCD80* (B), *PIGR* (C), and *PBHMT* (D) in RNA-seq data from 81 human SCLC tumors. **E.** Apoptosis analysis of KP11 SCLC cells treated with IL-6 blocking antibody in the presence or absence of TILs for 48 hours (n=4-6). Statistical significance was assessed by a t-test; error bars represent SD. **F.** Quantification for *Cd7*4 mRNA levels relative to *Gapdh* in KP11 cells co-culture with TILs, with or without IL-6-blocking antibodies (n=3). Statistical significance was assessed using a t-test; error bars represent s.e.m. **G.** Quantification of CD74 staining normalized to DAPI (n=8 mice, one tumor per mouse). Statistical significance was assessed using a t-test; error bars represent s.e.m. **H.** Representative immunofluorescence staining of CD74 (red) in KP1 SCLC allografts from control (Ctrl) and T cell-depleted (Depl.) mice, as in (G). DAPI stains DNA in blue. Scale bar, 50 µm. **I.** Representative immunofluorescence staining of CD3 (green) and CD74 (red) in KP1 SCLC allografts from control tumors (Ctrl) and tumors mixed with TILs cells (+TILs). DAPI stains DNA in blue. Scale bar, 50 µm. **J.** Quantification of CD3 staining as in (I), normalized to DAPI (n=7 mice, n=1-2 tumors per mouse). Statistical significance was assessed using a t-test; error bars represent s.e.m. **K.** Quantification of CD74 as in (I), normalized to DAPI (n=7 mice, n=1-2 tumors per mouse). Statistical significance was assessed using a t-test; error bars represent s.e.m. **L.** Representative immunofluorescence staining for CD3 (green) and CD74 (red) in KP11 SCLC allografts from control tumors (Ctrl) and tumors mixed with TILs cells (+TILs). DAPI stains DNA in blue. Scale bar, 50 µm. **M.** Quantification of CD3 staining as in (L), normalized to DAPI (n=7-9 mice, one tumor per mouse). Statistical significance was assessed using a t-test; error bars represent s.e.m. **N.** Quantification of CD74 staining as in (L), normalized to DAPI (n=7-9 mice, one tumor per mouse). Statistical significance was assessed using a t-test; error bars represent s.e.m. **O.** Representative immunoblot and quantification of phosphorylated AKT (pAKT) relative to total AKT normalized to HSP90 in KP11 SCLC cells cultured alone or co-cultured with TILs or splenic T cells (T^Spleen^). Statistical significance was assessed by a t-test; error bars represent SD (n=3). **P.** *Cd74* and *Mif* expression in KP1 and TILs co-culture assay, as described in Fig. 4A. **Q.** *Cd74* and *Mif* expression in KP11 allograft tumors with or without TILs, as described in Fig. 4A. **R.** Cell viability assay (AlamarBlue) of KP11 SCLC cells treated with ISO-1 (100 µg/mL and 250 µg/mL) or DMSO control (Ctrl) for 4 days (n=3). Statistical analysis was performed using two-way ANOVA; error bars represent s.e.m. **S.** Cell death analysis of KP11 SCLC cells treated with ISO-1 (100 µg/mL) or DMSO control (Ctrl), with or without TILs cells for 48 hours (n=4-6). Statistical significance was assessed using a t-test; error bars represent SD.

**Supplementary Figure S5, related to Figure 4:**
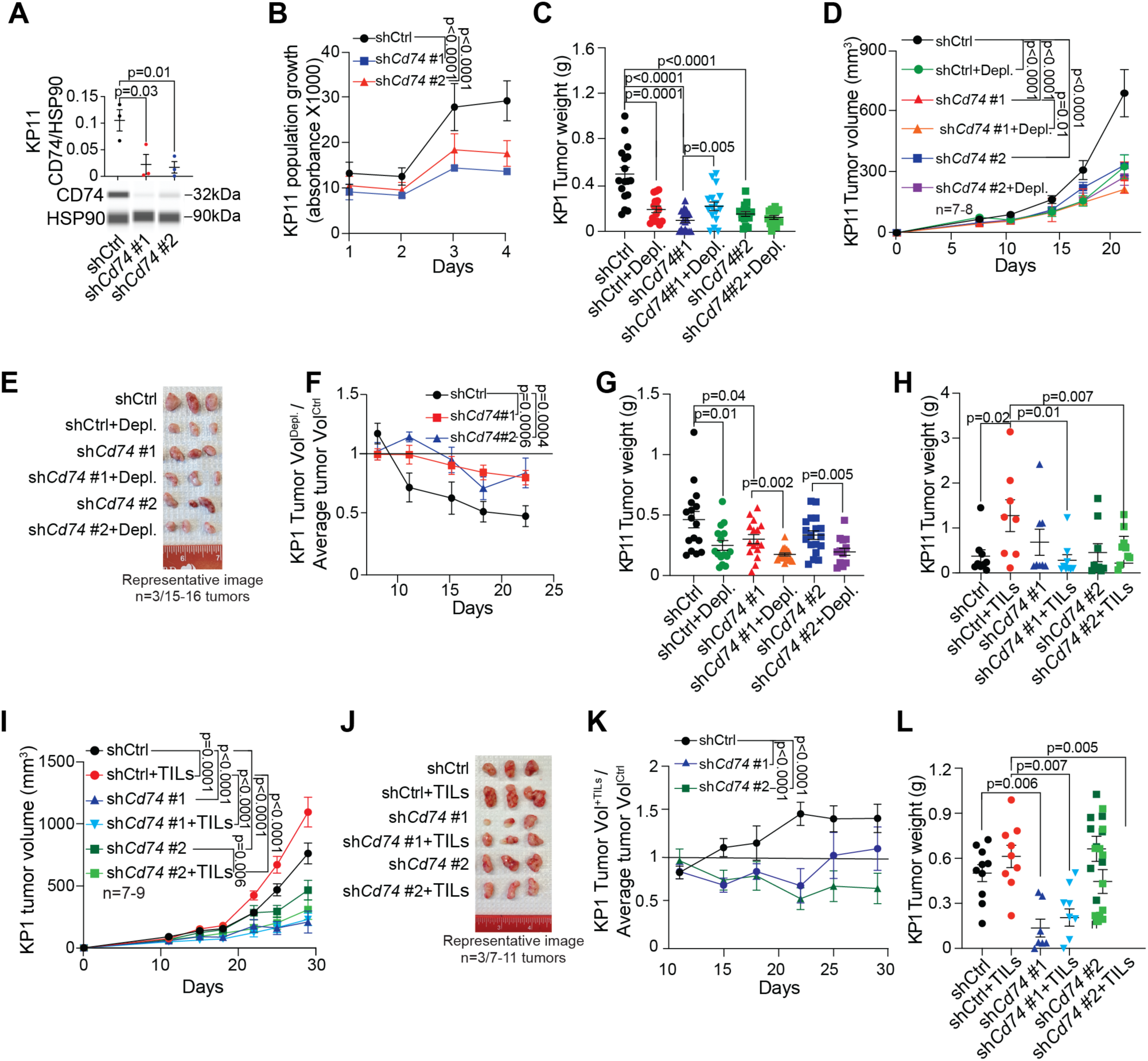
Expression of the pro-survival factor CD74 is induced in SCLC cells by TILs. **A.** Representative immunoassay and quantification of CD74 protein levels relative to HSP90 in control (shCtrl) and CD74 knock-down (sh*Cd74* #1 and #2) KP11 cells (n=3). Statistical significance was assessed using a t-test; error bars represent SD. **B.** Cell viability assay (AlamarBlue) of control (shCtrl) and *Cd74* knock-down (sh*Cd74* #1 and #2) KP11 cells for 4 days (n=3). Statistical analysis was performed using two-way ANOVA; error bars represent s.e.m. **C.** Tumor weight of KP1 SCLC allografts at terminal time point as in Fig. 4N (n=8-9 mice, 2 tumors per mouse, 1 experiment). Statistical significance was assessed using a t-test; error bars represent s.e.m. **D.** Tumor growth curves of KP11 SCLC allografts (n=7-8 mice, 2 tumors per mouse, 1 experiment) in control (shCtrl) and T cell-depleted mice (Depl.), with or without *Cd74* knock-down. Statistical analysis was performed using two-way ANOVA; error bars represent s.e.m. **E.** Representative images of KP11 SCLC allografts at terminal time point as in (D). **F.** Fold change in tumor size as in (D). Depletion ratio score calculated by dividing tumor volume at each time point by the mean volume of control tumors. Ratio >1 indicates an anti-tumor effect of T cells, ratio = 1 indicates no effect, and ratio <1 indicates a pro-tumor effect. In controls, ratio decreases (<1), whereas in CD74 knock-down tumors, ratio remains ∼1. Statistical analysis was performed using two-way ANOVA; error bars represent s.e.m. **G.** Tumor weight of KP11 SCLC allografts at terminal time point as in (D). Statistical significance was assessed using a t-test; error bars represent s.e.m. **H.** Tumor weight of KP11 SCLC allografts at terminal time point as in Fig. 4Q. Statistical significance was assessed using a t-test; error bars represent s.e.m. **I.** Tumor growth curves of control KP1 SCLC allografts and allografts co-injected with TILs cells (n=7-9 mice, 1-2 tumors per mouse, 1 experiment), with or without *Cd74* knock-down. Statistical analysis was performed using two-way ANOVA; error bars represent s.e.m. **J.** Representative images of KP1 SCLC allografts at terminal time point, as in (I). **K.** Fold change in tumor size as in (I). Tumor volume at each time point was divided by the mean volume of control tumors. As expected, the ratio increases for control tumors (TILs promote tumor growth), whereas it remains closer to 1 for *Cd74* knock-down tumors. Statistical analysis was performed using two-way ANOVA; error bars represent s.e.m. **L.** Tumor weight of KP1 SCLC allografts at terminal time point, as in (I). Statistical significance was assessed using a t-test; error bars represent s.e.m.

**Supplementary Figure S6, related to Figure 5:**
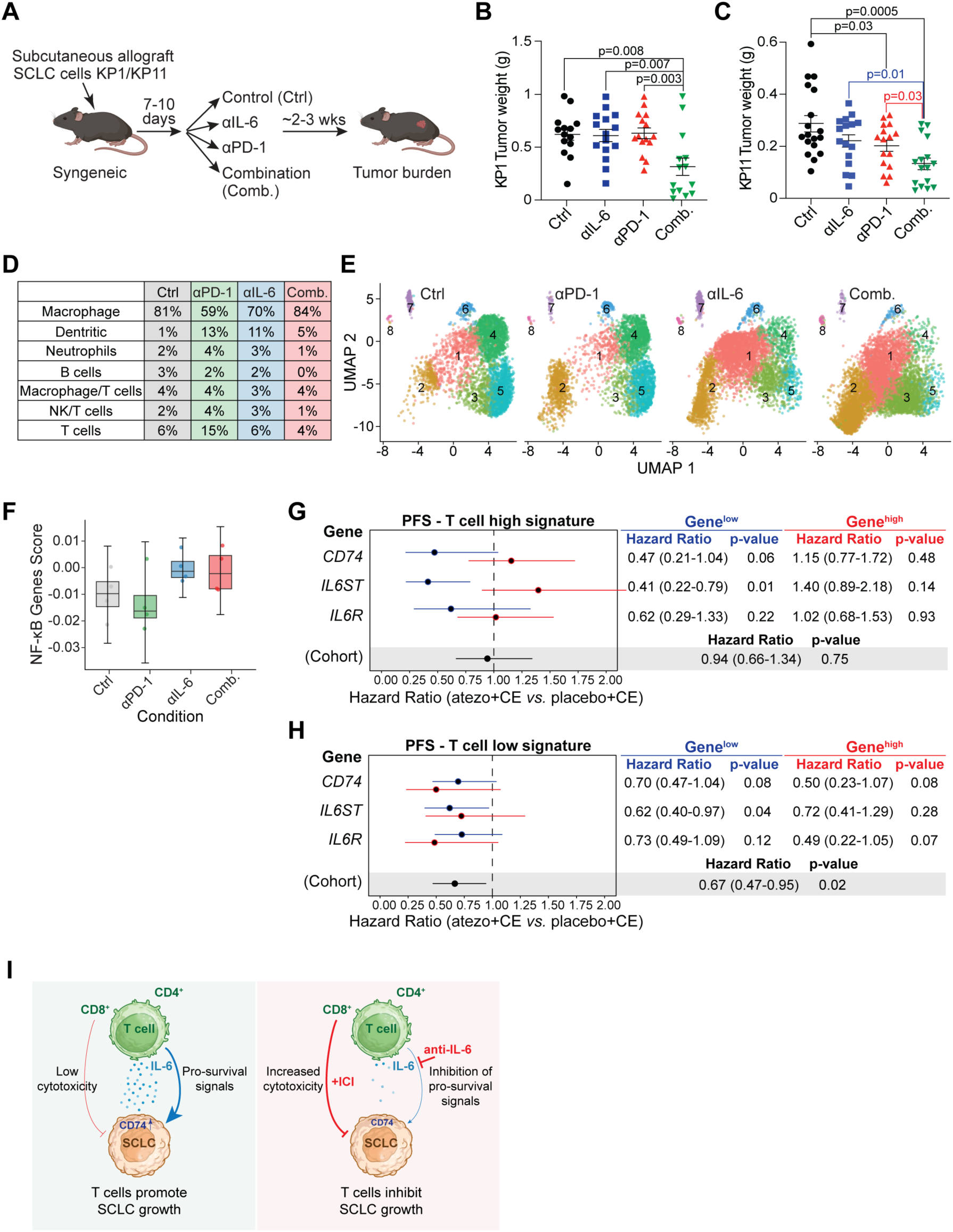
Combining IL-6 and PD-1 blockade inhibits SCLC growth. **A.** Cartoon of the approach used to test the combination therapy. **B.** Tumor weight of KP1 SCLC allografts at the terminal time point as in Fig. 5A. Statistical significance was assessed using a t-test; error bars represent s.e.m. **C.** Tumor weight of KP11 SCLC allografts at the terminal time point as in Fig. 5C. Statistical significance was assessed using a t-test; error bars represent s.e.m. **D.** Table showing the percentage of each immune cell cluster in each treatment condition as in Fig. 5F. **E.** Uniform Manifold Approximation and Projection (UMAP) plots of scRNA-seq data from SCLC cells isolated from KP11 allografts (n = 4 per treatment), comparing untreated, PD-1 blockade, IL-6 blockade, and combined treatments. Cell clusters are color-coded by population, illustrating changes in intratumoral SCLC heterogeneity. **F.** Box plot of NF-κB pathway scores in SCLC cells across the four experimental conditions. Statistical analysis by t-test; error bars represent SD. **G.** Forest plots showing progression-free survival (PFS) hazard ratios with 95% confidence intervals (CI) for atezolizumab plus carboplatin/etoposide (atezo+CE) versus placebo plus CE (placebo+CE) in the T signature > median cohort, stratified by gene expression. Low gene expression (≤ median) cohort shown in blue (Gene^low^), high gene expression (> median) cohort shown in red (Gene^high^). “Cohort” refers to the T signature > median without further subsetting. **H.** As in (G) for the T signature ≤ median cohort, stratified by gene expression. **I.** Model: (Left) In most cases of SCLC, T cells fail to control SCLC tumor growth, in part because of low cytotoxic activity and also because IL-6 produced by T cells can activate pro-survival signals in SCLC cells, including via induction of CD74. (Right) Combining immune checkpoint inhibition (ICI) to activate the cytotoxic activity of T cells and IL-6 blockade to suppress pro-survival signals may help inhibit SCLC growth.

